# Excitable axonal domains adapt to sensory deprivation in the olfactory system

**DOI:** 10.1101/2021.01.25.428132

**Authors:** Nicholas M George, Wendy B Macklin, Diego Restrepo

## Abstract

The axon initial segment, nodes of Ranvier, and the oligodendrocyte-derived myelin sheath have significant influence on the firing patterns of neurons and the faithful, coordinated transmission of action potentials to downstream brain regions. In the olfactory bulb, olfactory discrimination tasks lead to adaptive changes in cell firing patterns, and the output signals must reliably travel large distances to other brain regions along highly myelinated tracts. Whether myelinated axons adapt to facilitate olfactory sensory processing is unknown. Here, we investigate the morphology and physiology of mitral cell axons in the adult olfactory system, and show that unilateral sensory deprivation causes system-wide adaptations in axons. Mitral cell spiking patterns and action potentials also adapted to sensory deprivation. Strikingly, both axonal morphology and mitral cell physiology were altered on both the deprived and non-deprived sides, indicating system level adaptations to reduced sensory input. Our work demonstrates a previously unstudied mechanism of plasticity in the olfactory system.

## Introduction

In the olfactory bulb (OB), the firing rates and synchronized firing patterns of mitral (MC) and tufted cells encode vital olfactory information such as odor identity and odor salience (whether an odor deserves attention) (***Friedrich et al., 2004***; ***Doucette et al., 2011***; ***Lepousez and Lledo, 2013***; ***Gire et al., 2013***). The precise signals generated in the mouse OB must faithfully travel several millimeters along myelinated axonal tracts before reaching other brain regions such as the piriform cortex (***Nagayama et al., 2010***; ***Imamura et al., 2020***; ***Chon et al., 2020***; ***Walz et al., 2006***; ***Schwob and Price, 1984***). Downstream neurons in the piriform cortex are very sensitive to synchronized input, often failing to fire if the latency of incoming action potentials (AP)s is greater than 10ms ***(Franks and Isaacson, 2006; Luna and Schoppa, 2008***; ***Bolding and Franks, 2018***). Despite the importance of reliable, synchronized olfactory signals and the large distances these signals must travel, little is known about the structure and function of the myelinated axons that govern signal reliability and timing in the olfactory system.

Excitable axonal domains such as the axon initial segment (AIS) and nodes of Ranvier, as well as the insulating myelin sheath, regulate cellular excitability, firing patterns, and conduction speed (***Kole, 2011***; ***Evans et al., 2015***; ***Castelfranco and Hartline, 2016***). The AIS is the portion of the axon close to the soma and is defined by a high density of voltage gated sodium channels necessary for initiating APs and defining features of their kinetics and shape (***Kole et al., 2007***; ***Palmer and Stuart, 2006; Kole et al., 2008***). On myelinated axons, the myelin sheath and axon form specialized domains called nodes of Ranvier. Nodes of Ranvier allow for fast saltatory conduction by regenerating APs as they travel along an axon (***Susuki et al., 2013***; ***Castelfranco and Hartline, 2016***). The myelin sheath itself is produced by oligodendrocytes in the central nervous system and provides electrical insulation on stretches of axon between nodes of Ranvier. These stretches of insulated axon, which are vital for fast saltatory conduction, also provide trophic and metabolic support to axons (***Castelfranco and Hartline, 2016***; ***Meyer et al., 2018***; ***Fünfschilling et al., 2012***). Importantly, the organization of myelinated axons is not static. Both *in vitro* and *in vivo*, the AIS changes position, ion channel composition, and length in response to activity deprivation and stimulation paradigms, with significant consequences for cellular excitability (***Kuba et al., 2010***; ***Grubb and Burrone, 2010***; ***Evans et al., 2015***; ***Jamann et al., 2021***). Nodes of Ranvier develop specific sizes and spacing to optimize AP timing in the auditory brainstem (***Ford et al., 2015***), while the myelin sheath and oligodendrocyte lineage cells respond to neuronal activity and change throughout life (***Hill et al., 2018***; ***Hughes et al., 2018***; ***Gibson et al., 2014***). Computational modeling and experimental studies highlight the importance of myelinated axons for synchronized firing, AP reliability, and optimizing AP conduction speed (***Arancibia-Cárcamo et al., 2017***; ***Pajevic et al., 2014***; ***Kim et al., 2013***). Whether myelinated axons adapt to facilitate olfactory sensory processing is unknown.

Here, we investigated the AIS, nodes of Ranvier, and myelin sheaths (together referred to as myelinated axons) in the mouse OB and lateral olfactory tract (LOT). To determine whether these structures adapt to changing sensory input *in vivo*, we used adult unilateral naris occlusion (UNO) (***Baker et al., 1993***; ***Kass et al., 2013***; ***Coppola, 2012***) to suppress olfactory input and evaluated changes in myelinated axons and oligodendrocyte lineage cells. We found morphological adaptations in the AIS and nodes of Ranvier, and physiological changes in AP generation in MCs. In contrast to developmental UNO (***Collins et al., 2018***), we found no changes in relative myelin sheath thickness or oligodendrocyte lineage cell density. Whole cell patch-clamp experiments on MCs revealed increased AP amplitude and width following UNO, as well as altered cell firing. Our findings raise the possibility that excitable axonal domains may adapt to facilitate olfactory sensory processing in the adult.

## Results

### Quantification of mitral cell axon initial segments in the olfactory bulb and nodes of Ranvier in the lateral olfactory tract

The coordinated, precise firing of groups of MCs encodes olfactory information in the OB, and firing patterns of MCs change as an animal learns to identify odors in olfactory discrimination tasks (***Gire et al., 2013***; ***Doucette et al., 2011***; ***Lepousez and Lledo, 2013***; ***Chu et al., 2016***; ***Gschwend et al., 2015***). The AIS is the site of AP initiation, and its morphological structure and ion channel composition determines AP threshold, AP width and amplitude, and other important firing properties of the cell (***Palmer and Stuart, 2006***; ***Kole et al., 2007**,**2008***; ***Kole and Stuart, 2012***; ***Jamann et al., 2021***).

MCs are known to have identifiable AISs similar to other brain regions ***(Lorincz and Nusser, 2008; Price and Powell, 1970***; ***Hinds and Ruffett, 1973***), but their structure and size distributions in OB are not well understood. We measured the length of MC AISs in 3D volumes using immunohistochemistry and confocal microscopy (***Figure 1***). The OB is a highly stratified structure whose layers are clearly delimited using the nuclear label Hoechst (***Figure 1***B). The histochemical dye Nissl (NeuroTrace, Thermo Fisher Scientific) broadly labels cells in the OB, and MCs could be clearly identified based on their large Nissl+ somas and their location in the mitral cell layer (MCL). AnkyrinG (AnkG) is a cytoskeletal scaffolding protein essential for the structure and function of the AIS ***(Hedstrom et al., 2008***). The AnkG+AIS of MCs extended directly off of the MC soma, often immediately into the first myelinated internode (***Figure 1***B’, white arrowheads mark AIS start, yellow mark the end). AISs were measured in 3D volumes using the ImageJ/Fiji plugin Simple Neurite Tracer (SNT) (***Arshadi et al., 2020***), starting at the AnkG label originating at the base of the Nissl+ cell body and terminating at the distal end of the AnkG label. The mean length of MC AISs in the OB was 25.7 ± 3.81 μm (lengths are given as mean ± standard deviation [SD], *n* = 687 AISs from 4 animals [***Figure 1C***]).

**Figure 1.**
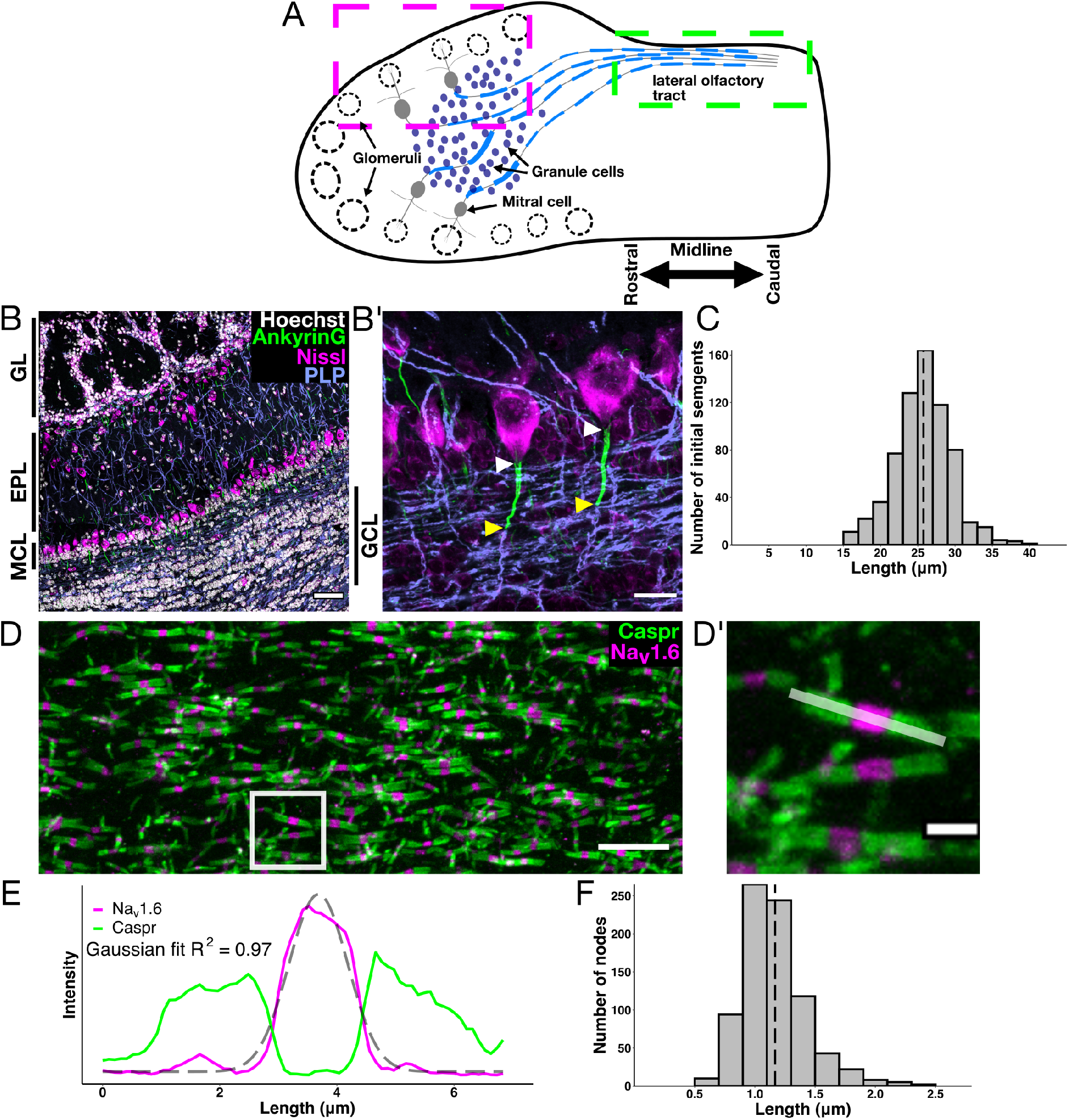
Characterization of excitable axonal domains in the olfactory system. (**A**) Diagram of a transverse section of the olfactory bulb (OB) and rostral lateral olfactory tract (LOT). Boxes indicate the areas where measurements were performed for the AIS and nodes of Ranvier. Magenta box (**A**) corresponds to an example region for AIS length measurements (**B** and **B’**), and the green boxed region (**A**) corresponds to an example region for node of Ranvier measurements (**D**, **D’**). (**B**) Immunohistochemistry for nuclei (Hoechst, white), the axon initial segment (AIS) (AnkG, green), cell bodies (Nissl, magenta), and myelin (proteolipid protein, PLP, blue). The functional layers of the OB are labeled on the edges of the image (GL = glomerular layer, EPL = external plexiform layer, MCL = mitral cell layer, GCL = granule cell layer). Scale bar in (**B**) is 50μm. (B’) Expanded region highlighting Nissl labeling of MC bodies and AnkG+ AISs in the MCL. White arrowheads label the start of MC AISs and yellow arrowhead show the termination. Scale bar in (**B’**) is 15μm. (**C**) Distribution of AIS lengths in the adult OB. Dashed line indicates mean length. (**D**) Nodes of Ranvier in the LOT corresponding to the region highlighted in the green box in (**A**). Nav1.6 (magenta) marks the excitable nodal region, Caspr (green) marks flanking paranodal junctions. Scale bar in (**D**) is 10μm. (**D’**) Expanded node of Ranvier from white box in (**D**). Scale bar in (**D’**) is 2μm. (**E**) Fluorescence intensity profile of the node of Ranvier in (D’, light white line) with a Gaussian fit of Nav1.6 (grey dashed line). The Gaussian fit of Nav1.6 was used to measure node length (see Methods). (**F**) Distribution of node lengths in the LOT. Dashed line indicates mean length.

While precise firing patterns are generated in the OB, MC axons must carry these signals out of the OB to the piriform cortex and other brain regions via the highly myelinated LOT ***(Nagayama et al., 2010; Imamura et al., 2020***; ***Chon et al., 2020***; ***Walz et al., 2006***; ***Schwob and Price, 1984***). Nodes of Ranvier are the axonal gaps between the myelin sheaths along the axon. Nodes of Ranvier regenerate the AP as it travels along an axon in a process called saltatory conduction ***(Susuki et al., 2013***). Tuning of node of Ranvier length or channel composition may be a sensitive way to regulate AP conduction velocity and synchrony (***Arancibia-Cárcamo et al., 2017***). To assess the structure of nodes of Ranvier in the LOT, we used antibodies directed against contactin-associated protein (Caspr) and the voltage gated sodium channel Na_v_1.6. Caspr marks the site of myelin sheath to axon adhesion and serves as an important ion diffusion barrier (***Bhat et al., 2001***; ***Rios et al., 2003***), while Na_v_1.6 is the primary voltage gated sodium channel in mature nodes, responsible for regenerating the AP (***Caldwell et al., 2000***; ***Boiko et al., 2001***; ***Rasband et al., 2003***). The LOT contains a high density of nodes clearly identified by Na_v_1.6 flanked by Caspr (***Figure 1***D). We measured node of Ranvier length manually by fitting a Gaussian to the fluorescence intensity profile of antibody labeled Na_v_1.6 and measuring the full width at half maximum of that fit (***Figure 1***D’,E; see Methods). The mean length of nodes of Ranvier in the LOT was 1.17 ± 0.267μm (mean ± SD *n* = 811 nodes from 4 animals) (***Figure 1***F).

### Quantification of oligodendrocyte lineage cells and the myelin sheath

In the central nervous system, the myelin sheath is made by oligodendrocytes. Oligodendrocytes develop from oligodendrocyte progenitor cells (OPCs), a population long known to proliferate, differentiate into oligodendrocytes, and produce myelin in response to neuronal activity ***(Barres and Raff, 1993; Gibson et al., 2014***; ***McKenzie et al., 2014***; ***Young et al., 2013***). In the mouse olfactory system, a significant period of OPC differentiation and myelination begins around postnatal day (P) 10 and is largely complete by P30 (***Philpot et al., 1995***; ***Collins et al., 2018***). Little is known about oligodendrocyte lineage cells in the mature olfactory system. The transcription factor Olig2 marks the entire oligodendrocyte lineage, while PDGFR*α* marks immature OPCs and CC1 marks mature oligodendrocytes. We counted OPCs using antibodies directed against Olig2 and PDGFR*α*, and oligodendrocytes using antibodies against Olig2 and CC1. We counted oligodendrocytes and OPCs in the granule cell layer (GCL) and the LOT of adult animals in 3D confocal volumes (***Figure 2A***-D). The GCL contained an average of 6.43e-6 ± 3.17e-6 OPCs/μm^3^ and 3.36e-5 ± 5.53e-6 oligodendrocytes/μm^3^ (cell counts are given as mean density ± SD). The LOT contained 7.49e-6 ± 2.1e-6 OPCs/μm^3^ and 9.98e-5 ± 7.98e-6 oligodendrocytes/μm^3^ (all cell counts were performed with a systematic random sampling scheme using 3-5 sections per animal, where *n* animals *≥* 4; see Methods).

**Figure 2.**
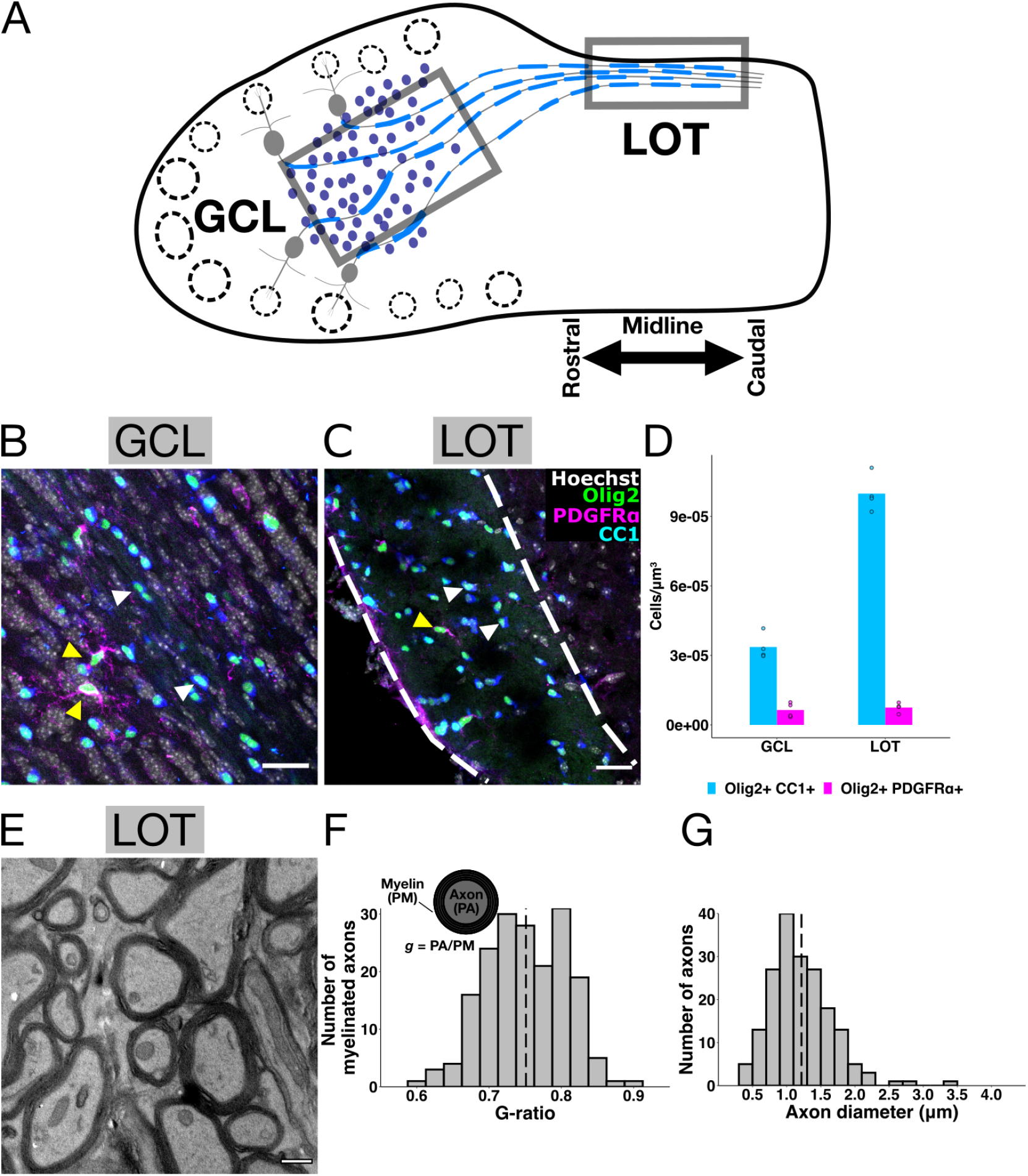
Characterization of oligodendrocyte lineage cells and myelin sheaths in the OB and LOT. (**A**) Diagram ofatransverse section of the OB and LOT with grey boxes indicating the GCL and LOT. (**B**) Oligodendrocyte lineage cells in the GCL. (**C**) Oligodendrocyte lineage cells in LOT (area between dashed white lines). Yellow arrows indicate oligodendrocyte progenitor cells (OPCs) and white arrows indicate oligodendrocytes. Scale bars in (**B** and **C**) are 25μm. (**D**) Oligodendrocyte lineage cell quantification in GCL and LOT. Points indicate individual animals (*n* = 4 animals) and bars indicate the group means. (**E**) Electron micrograph (EM) image of myelinated axons in a coronal section of LOT. Scale bar in (**E**) is 0.5μm. (**F**) Diagram demonstrating *g*-ratio calculations and histogram of *g*-ratio distribution in LOT. To calculate*g*-ratio, the perimeter of the axon (PA) was divided bythe perimeter of the surrounding myelin sheath (PM) to yield *g*. Histogram shows distribution of *g*-ratios in LOT. Dashed line histogram indicates mean *g*-ratio. (**G**) Distribution of axon diameters in LOT. Dashed line indicates mean axon diameter.

The myelin sheath provides electrical insulation and vital trophic/metabolic support to axons (***Castelfranco and Hartline, 2016***; ***Meyer et al., 2018***; ***Fünfschilling et al., 2012***). The *g*-ratio (presented as axon perimeter / total fiber perimeter), is a measure of myelin thickness relative to axon diameter. The *g*-ratio has long been used in computational modeling studies as a parameter to assess the conduction velocity along myelinated axons (***Rushton, 1951***; ***Chomiak and Hu, 2009***). Since axon size and myelin thickness (*g*-ratio) have significant effects on axonal conduction velocity and AP fidelity (***Chomiak and Hu, 2009***; ***Kim et al., 2013***; ***Etxeberria et al., 2016***), we measured the *g*-ratio of myelinated axons in the LOT between 2.45-3.05mm anterior of the bregma suture (***Figure 2**D-H*). The mean *g*-ratio was 0.751 ± 0.0549, while mean axon diameter was 1.22 ± 0.456μm (***Figure 2***F,H) (*n* = 184 axons from 2 animals).

### Unilateral naris occlusion causes adaptations in mitral cell axon initial segments and nodes of Ranvier

The OB is a remarkably plastic structure. New neurons are incorporated into functional circuits in the OB throughout life (***Whitman and Greer, 2007***; ***Yamaguchi et al., 2013***; ***Lazarini et al., 2009***), and the firing patterns of MCs are known to change significantly as an animal learns an olfactory discrimination task (***Losacco et al., 2020***; ***Chu et al., 2016***; ***Friedrich et al., 2004***; ***Doucette et al., 2011; Lepousez and Lledo, 2013***; ***Gire et al., 2013***; ***Gschwend et al., 2015***). Little is known about whether axonal domains adapt to changing olfactory input in adult animals. To test whether myelinated axons respond to changing olfactory input in adult mice, we performed UNO on P60 mice for 30 days and measured the length of MC AISs and nodes of Ranvier in the LOT.

UNO is a model for sensory deprivation in which one naris is surgically closed to block sensory input. One of the hallmarks of UNO is an activity-dependent decrease in tyrosine hydroxylase (TH) mRNA and protein expression in a subset of dopaminergic periglomerular cells in the glomerular layer (GL) (***Sawada et al., 2011***; ***Baker et al., 1993***). To assess the efficiency of UNO, we measured the fluorescence intensity of TH in the GL of P60 mice that underwent UNO for 30 days (called Naris Occlusion) and control animals (cage mate sham surgery, called Control) (***Figure 3***). Within Naris Occlusion animals, we noted a significant decrease (~45%) of TH fluorescence intensity in the Occluded bulb GL compared to the Open bulb GL (paired *t*-test, *t(6)* = 9.58, *p* = 7.4e-5, *n* = 7 animals). Control animals showed no significant difference in TH intensity between Left and Right bulbs (***Figure 3***D, paired *t*-test, *t*(3) = −0.383, *p* = 0.727, *n* = 4 animals). We compared the relative intensity (Occluded side/Open side for Naris Occlusion animals, and Left side / Right side for Control animals) and found that the relative intensity was roughly equal (centers on 1) in Control animals, but was significantly reduced in Naris Occlusion animals, indicating effective UNO (***Figure 3***E, *t*-test, *t*(7.2)= 6.25, *p* = 0.00038).

**Figure 3.**
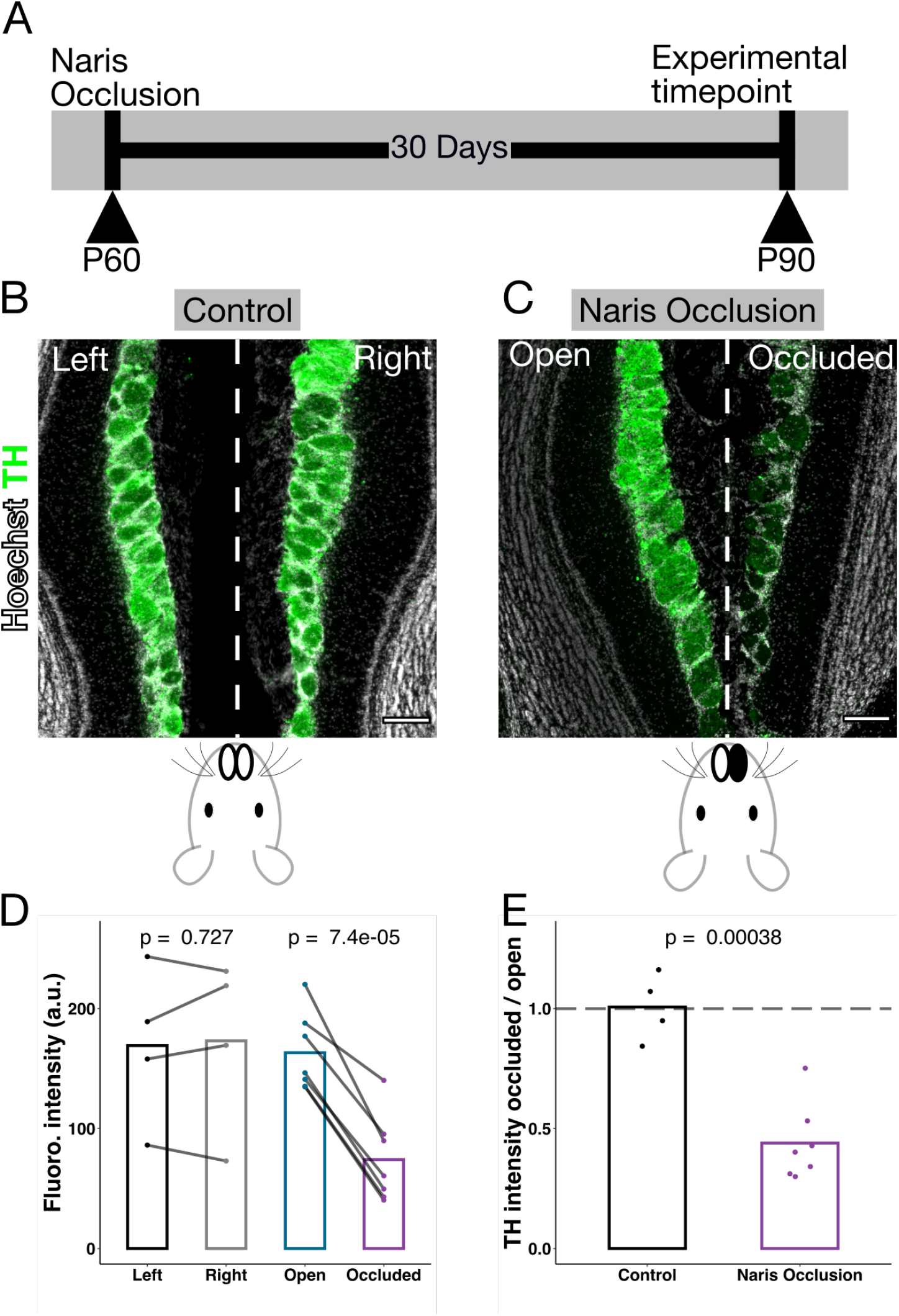
Unilateral naris occlusion experimental design. (**A**) Experimental timeline. Postnatal day (P) 60 animals underwent unilateral naris occlusion (UNO) for 30 days and were analyzed at P90. (**B**) Example bilateral horizontal section ofOB showingTH immunolabeling in a P90 Control animal. (**C**) Example bilateral horizontal section ofOB showingTH immunolabeling in a P90 animal following 30 days UNO (Naris Occlusion). Scale bars in **B** and **C** are 150μm. (**D**) Fluorescence intensity is not significantly different between the Left and Right sides of Control animals (paired *t*-test, *t*(3) = −0.383, *n* = 4 animals). There is a significant decrease in fluorescence intensity between the Open and Occluded bulbs of Naris Occlusion animals (paired *t*-test, *t*(6) = 9.58, *n* = 7 animals). Intensity is given in arbitrary units (a.u.). (**E**) Plot showing relative intensity of OBswithin Control animals orwithin Naris Occlusion animals. Naris Occlusion animals have significantly lower relative TH intensity compared to Control animals. Dashed line at 1 indicates equal intensity on either side (Right / Left for Control or Occluded / Open for Naris Occlusion). Relative TH intensity was significantly reduced in Naris Occlusion animals compared to Control (*t*-test, *t*(7.2) = 6.25).

Do MC AISs adapt to changing sensory input following UNO? We measured MC AISs in 3D volumes (as in ***Figure 1***, see Methods) in both 30 day Naris Occlusion animals and Controls. The distribution of AIS lengths was not significantly different between the Left and Right sides of Control animals (Kolmogrov-Smirnov (KS) test, *D* = 0.0599, *p* = 0.592, *n* = 678 AISs from 4 animals), but in Naris Occlusion animals, the Open and Occluded sides were significantly different (KS test, *D* = 0.128, *p* = 3.97e-7, *n* = 1,894 AISs from 7 animals) (***Figure 4***A-C).

**Figure 4.**
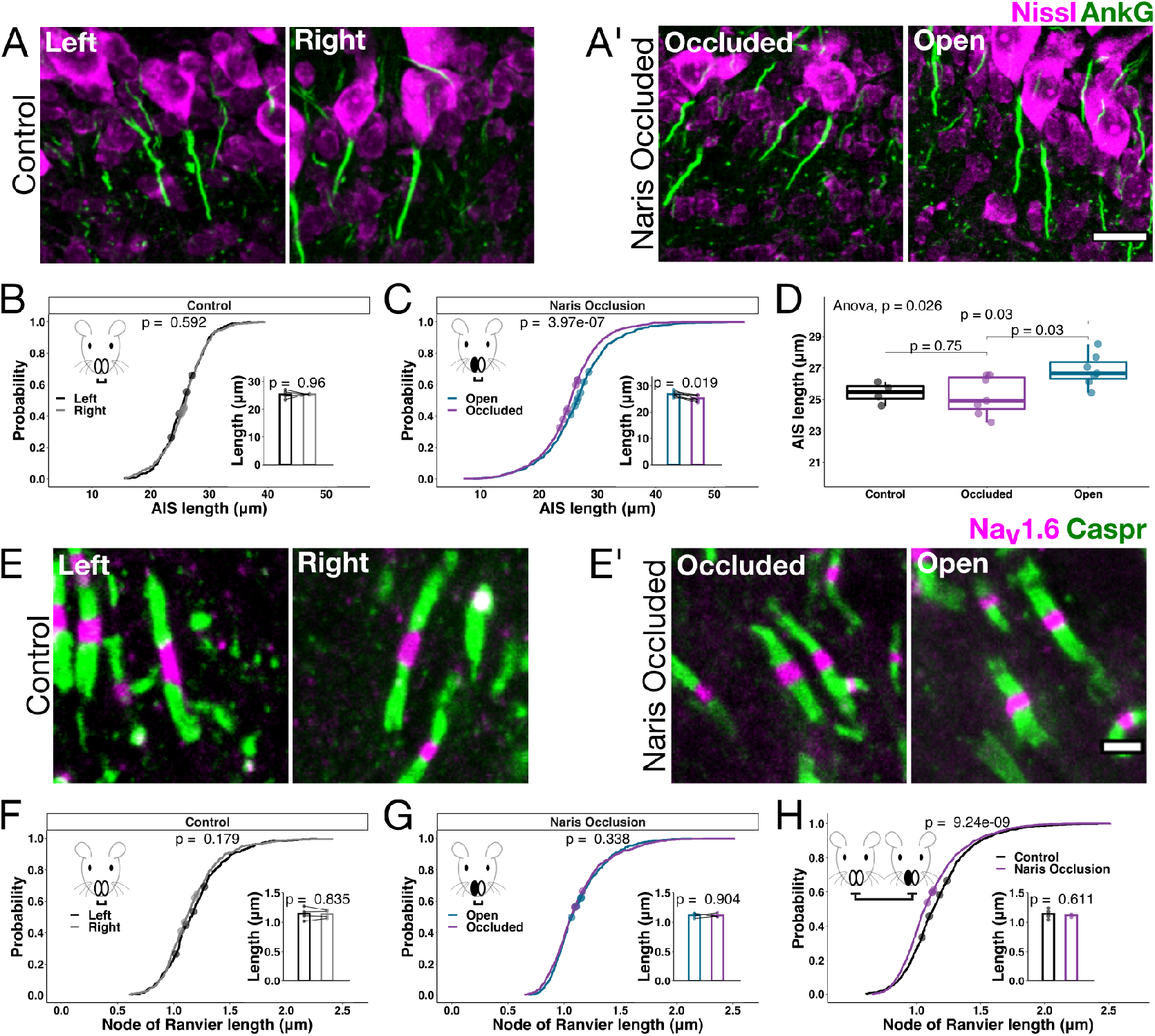
MC axonal adaptations following 30 days of UNO. (**A**) Example MC AISs from the left and right OB of Control animals. (**A’**) Example MC AISs from the Occluded and Open OB from UNO animals. Scale bar in (**A’**) and (**A**) and is 15μm. (**B**) Empirical distributions comparing the AIS lengths from the left and right OBs of Control animals. *P*-value is from a KS-test comparing the distributions (KS-test *D* = 0.0599, *n* = 678 AISs from 4 animals). Points along the empirical distribution show the mean length of each OB’s AISs mapped to the probability within the respective distribution. Insert shows within-animal comparison of mean AIS lengths. Points in the bar plot inserts represent the mean AIS length from an animal’s left or right OB, and the bars represent the mean value for that group (paired *t*-test, *t*(3) = −0.0548). *N* = 678 AISs from 4 animals. (**C**) Similar to (**B**) but comparing the empirical distribution of AISs from Open and Occluded OBs from UNO animals (KS-test *D* = 0.128, *n* = 1,894 AISs from 7 animals). Insert shows within-animal comparison of mean AIS lengths in Open vs. Occluded bulbs from Naris Occlusion animals (as in (**B**), paired *t*-test, *t*(6) = 3.18). (**D**) Mean AIS lengths calculated on a per-animal OB basis (Control combines the left and right OBs from Control animals, while Naris Occlusion Occluded and Open OBs were kept separate). AIS lengths from Control OBs were not significantly different from AIS lengths from Occluded OBs (*t*-test *t*(9) = 0.32), but AISs from Open OBs were significantly longer than both Control (*t*-test, *t*(8.66) = −2.84) and Occluded OBs (*t*-test, *t*(11.6) = −2.69). False discovery rate (FDR) corrected *p* values are shown in plot. (**E**) Example nodes of Ranvier in the LOT from the Left and Right sides of Control animals. (**E’**) Similar to (**E**), but in UNO animals. Scale bar in (**E’**) and (**E**) and is 2μm. (**F**) Empirical distributions comparing node of Ranvier lengths from the left and right LOTs of Control animals. *P*-value is from a KS-test comparing node of Ranvier length distributions (KS-test *D* = 0.0773, *n* = 811 nodes from 4 animals), displayed similar to (**B-C**). Insert shows within-animal between side comparison of mean node of Ranvier lengths. Points represent the mean node of Ranvier length from an animal’s left or right LOT, and the bars in the insert represent the mean value for that side (paired *t*-test, *t*(3) = 0.227). (**G**) Similar to (**F**), but comparing node of Ranvier lengths from the Open and Occluded LOTs of Naris Occlusion animals (KS-test *D* = 0.0508, *n* = 1,409 nodes from 4 animals). Insert similar to (**F**), showing within-animal between side comparison of mean node of Ranvier lengths (paired *t*-test, *t*(3) = 0.131). (**H**) Comparison of node of Ranvier length distributions for Control and Naris Occlusion. *P*-value is from a KS-test comparing the node of Ranvier lengths for LOTs from Control animals or Naris Occlusion animals (KS-test *D* = 0.137). Points along the distribution show individual animal means mapped to the distribution. Insert shows a per-animal comparison of mean node length (Control vs. Naris Occlusion). The points represent per-animal means and bars represent the group means (*t*-test, *t*(3.38) = 0.56).

For each group (Control and Naris Occlusion), we summarized the data within animals and performed paired *t*-tests (inserts, ***Figure 4***B,C). The Left and Right sides of Control animals were not significantly different (paired *t*-test, *t*(3) = −0.0548, *p* = 0.96, *n* = 4 animals), but the Open and Occluded sides of Naris Occlusion animals were significantly different (paired *t*-test, *t*(6) = 3.18, *p* = 0.019, *n* = 7 animals).

Strikingly, Naris Occlusion animals (Open and Occluded sides combined) had a larger range of AIS lengths than Control animals (7.06 - 55.1 μm, SD = 5.83μm for Naris Occlusion, 15.4 - 39.5μm, SD = 3.81 μm for Control). Indeed, when we compared the variances of the two main distributions (Control vs. Naris Occlusion), they were significantly different (Fligner-Killeen test *χ*^2^(1) = 59.5, *p* = 1.26e-14). Within the Control group (Left vs. Right), we found no statistically significant difference in variance (Fligner-Killeen test *χ*^2^(1) = 0.0106, *p* = 0.918), indicating homogeneous variance between Left and Right sides of Control animals. However, within Naris Occlusion animals (Open vs. Occluded), variance was significantly different (Fligner-Killeen test *χ*^2^(1) = 9.97, *p* = 0.00159). A larger variance of AIS lengths in UNO animals could have implications for information encoding within the OB by increasing the diversity of AP shapes and firing frequencies from MCs (see Discussion).

To further investigate the difference in length distributions between Control and Naris Occlusion animals, we calculated the mean AIS length per animal and bulb. Given the consistency of AIS lengths in Control animals, we combined the Left and Right bulbs into one group. AIS lengths from Control bulbs, Open bulbs, and Occluded bulbs were significantly different from one another (***Figure 4***D, ANOVA, *F*(2,15) = 4.72, *p* = 0.026). Surprisingly, AISs from Open bulbs were significantly longer than Control bulbs (*t*-test, *t*(8.66) = −2.84, False discovery rate corrected [FDR] *p* = 0.03, mean length Open = 26.9 ± 1.02μm, mean length Control = 25.4 ± 1.04μm), and Occluded bulbs (*t*-test, *t*(11.6) = −2.69, FDR *p* = 0.03, mean length Open = 26.9 ± 1.02μm, mean length Occluded = 25.2 ± 1.24μm). Interestingly, Occluded bulbs were not significantly different from Control bulbs (*t*-test *t*(9) = 0.322, FDR *p* = 0.75, mean length Occluded = 25.2 ± 1.24μm, mean length Control = 25.4 ± 1.04μm).

Together, our data indicates that 30 days of UNO causes both a significant increase in MC AIS length in the Open bulb compared to Control and Occluded bulbs, and increases the variance of the entire distribution of AIS sizes in UNO animals relative to Controls.

Changes in (or loss of) nodes of Ranvier are predicted to have significant effects on conduction velocity and AP reliability (***Hamada et al., 2017***; ***Arancibia-Cárcamo et al., 2017***; ***Kim et al., 2013***), both of which are vital for olfactory sensory processing in downstream brain regions such as the piriform cortex (***Franks and Isaacson, 2006***; ***Luna and Schoppa, 2008***; ***Bolding and Franks, 2018***). To investigate whether nodes of Ranvier adapt to sensory deprivation in the olfactory system, we measured the length of nodes (as described in ***Figure 1***D’, E; see Methods) in Naris Occlusion animals and Controls. The distribution of node of Ranvier lengths between the Left and Right sides of Control animals was not significantly different (KS test, *D* = 0.0773, *p* = 0.179, *n* = 811 nodes from 4 animals). The distribution of node lengths between the Open and Occluded sides of Naris Occlusion animals was also not significantly different (KS test, *D* = 0.0508, *p* = 0.338, *n* = 1,409 nodes from 4 animals). Similarly, when the data was summarized by animal, we found no significant differences (***Figure 4***, inserts F and G, Control paired *t*-test, *t*(3) = 0.227, *p* = 0.835, Naris Occlusion paired *t*-test, *t*(3) =0.131, *p* = 0.904).

Since between bulb distributions (Left vs. Right and Open vs. Occluded) were not different in either group, we next compared the distribution of node of Ranvier lengths from Control animals to Naris Occlusion animals (***Figure 4***H). We found that the distribution of node lengths was significantly different between the Control and Naris Occlusion animals (KS test, *D* = 0.137, *p* = 9.24e-9). However, when we directly compared the mean values per animal, values were not significant (*t*- test, *t*(3.38) = 0.56, *p* = 0.611 [***Figure 4***H insert]). In the case of nodes of Ranvier, we found no evidence of differing variances between the Left and Right sides of Control animals (Fligner-Killeen tests, *χ*^2^(1) = 1.15, *p* = 0.283), the Open and Occluded sides of Naris Occlusion animals (Fligner-Killeen tests, *χ*^2^(1) = 0.564, *p* = 0.453), or Naris Occlusion vs. Control (Fligner-Killeen tests, *χ*^2^(1) = 2.95, *p* = 0.086). The lack of changes *within* animals but difference *between* treatment groups points to a system level adaptation to Naris Occlusion, affecting both Open and Occluded sides to a similar degree.

Together, these data point to a system-level adaptation in node of Ranvier and AIS length distributions following Naris Occlusion. Nodes of Ranvier are formed from axon-myelin interactions (***Susuki et al., 2013***), so we next investigated whether oligodendrocyte lineage cells or the myelin sheath adapt to adult UNO.

### Unilateral naris occlusion does not affect oligodendrocyte lineage cells or the myelin sheath

Do oligodendrocyte lineage cells and the myelin sheath respond to UNO in adults? We quantified OPC and oligodendrocyte density in Control and Naris Occlusion animals in the GCL and LOT after 30 days of UNO (***Figure 5***). We found no significant difference between the Left and Right sides of Control animals in the density of GCL oligodendrocytes (paired *t*-test, *t*(3) = −0.364, *p* = 0.74, *n* = 4 animals) or GCL OPCs (paired *t*-test, *t*(3) = 1.53, *p* = 0.223, *n* = 4 animals). We also found no significant difference between the Open and Occluded sides of Naris Occlusion animals in GCL oligodendrocytes (paired *t*-test, *t*(3) = 2.43, *p* = 0.0938, *n* = 4 animals) or GCL OPCs (paired *t*-test, *t*(3) = −0.516, *p* = 0.642, *n* = 4 animals), although three of the four animals had a reduction in oligodendrocytes in Occluded bulbs (***Figure 5***A). There was no significant difference between the main groups (Control and Naris Occlusion) in GCL oligodendrocytes (*t*-test, *t*(5.41) = 0.192, *p* = 0.855, *n* = 8 animals) or GCL OPCs (***Figure 5***, row A, *t*-test, *t*(5.22) = −0.668, *p* = 0.533, *n* = 4 animals/group).

**Figure 5.**
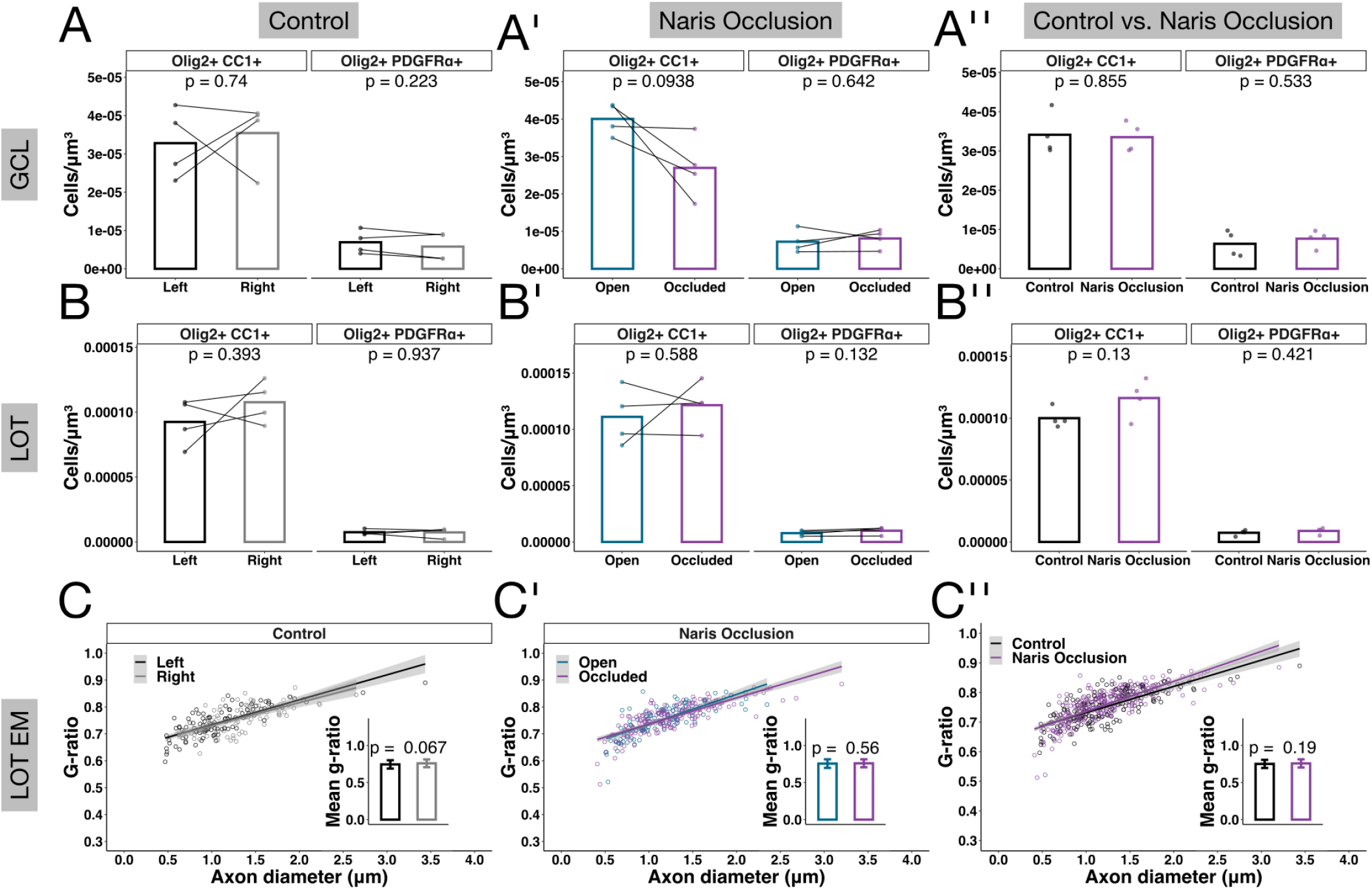
Oligodendrocyte lineage cells and myelinated axons after 30 days of UNO. (**A**) Comparison of GCL oligodendrocyte lineage cell density in the Left and Right bulbs of Control animals. We noted no significant differences between the Left and Right bulbs of Control animals in oligodendrocytes (paried *f*-test, *t(3)* = −0.364) or OPCs (paried *t*-test, *t*(3) = 1.53). Points represent within bulb means for each animal, *p*-values represent paired t-tests, *n* = 4 animals. (A’) Comparison of GCL oligodendrocyte lineage cell density in the Open and Occluded bulbs of Naris Occlusion animals. We found no significant difference in cell density between the Open or Occluded bulbs in oligodendrocytes (paried *t*-test, *t*(3) = 2.43) or OPCs (paried *t*-test, *t*(3) = 0.516). Points represent within bulb means for each animal, *p*-values represent paired *t*-tests, *n* = 4 animals. (**A”**) Comparison of LOT oligodendrocyte lineage cell density in Naris Occlusion vs. Control animals. Taken as a group, Control and Naris Occlusion animals had similar numbers of oligodendrocytes (*t*-test, *t*(5.41) = 0.192) and OPCs (*t*-test, *t*(5.22) = −0.668). Points represent per-animal means, *p*-values represent *t*-tests, *n* = 8 animals. (**B**) Comparison of LOT oligodendrocyte lineage cell density in the Left and Right bulbs of Control animals. We found no significant differences in oligodendrocyte (paried *t*-test, *t*(3) = −0.995) or OPC density (paried *t*-test, *t*(3) = 0.085) in LOT, quantification similar to (**A**), *n* = 4 animals. (**B’**) Comparison of LOT oligodendrocyte lineage cell density in Open and Occluded bulbs of Naris Occlusion animals. We found no significant differences in oligodendrocyte (paried *t*-test, *t*(3) = 0.606) or OPC density (paried *t*-test, *t*(3) = −2.05) in LOT, quantification similar to (**A’**), *n* = 4 animals. (**B”**) Comparison of LOT oligodendrocyte lineage cell density in Naris Occlusion vs. Control animals. Taken as a group, there was again no significant difference in oligodendrocyte lineage cell density between Naris Occlusion and Control animals (oligodendrocytes *t*-test, *t*(4.45) = −1.86, OPCs *t*-test, *t*(5.87) = 0.806). Quantification was similar to (**A”**), *n* = 8 animals. (**C**) Comparison of myelinated axons in the left and right LOTs of Control animals. There was no significant difference in mean *g*-ratio between the left and right sides of Control animals (*t*-test, *t*(179) = 1.85, *n* = 184 axons). Points in scatter plot represent individual axons, error bars in bar plot insert represent per-group standard deviation (SD). (**C’**) Comparison of myelinated axons in the Open and Occluded LOTs of Naris Occlusion animals. There was no significant difference in mean *g*-ratio between the Open and Occluded sides of Naris Occlusion animals (*t*-test, *t*(211) = −0.591). Data presented as in (**C**) *n* = 305 axons. (**C”**) Comparison of myelinated axons in Naris Occlusion vs. Control animals. Treated as a group, there was no significant difference between Naris Occlusion and Control animals (*t*-test, *t*(388) = −1.32).

Similarly, in the LOT, we found no significant differences between the Left and Right sides of Control animals in LOT oligodendrocytes (paired *t*-test, *t*(3) = −0.995, *p* = 0.393, *n* = 4 animals) or OPCs (paired *t*-test, *t*(3) = 0.085, *p* = 0.937, *n* = 4 animals). There were no significant differences between the Open and Occluded sides of Naris Occlusion animals in LOT oligodendrocytes (paired *t*-test, *t*(3) = −0.606, *p* = 0.588, *n* = 4 animals) or OPCs (paired *t*-test, *t*(3) = −2.05, *p* = 0.132, *n* = 4 animals). There was also no significant difference between the main groups (Control and Naris Occlusion) in LOT oligodendrocytes (***Figure 5***, row B, *t*-test, *t*(4.45) = −1.86, *p* = 0.13, *n* = 8 animals) or OPCs (*t*-test, *t*(5.87) = −0.866, *p* = 0.421, *n* = 4 animals/group).

While we found no change in the density of oligodendrocyte lineage cells, myelin sheaths themselves can undergo remodeling by pre-existing oligodendrocytes (***Auer et al., 2018***). To determine whether myelin sheaths changed in response to UNO, we quantified myelinated axons in the LOT of Naris Occlusion and Control animals by measuring the *g*-ratio (see Methods and ***Figure 2***F). We noted no significant differences between the Left and Right LOTs of Control animals (*t*-test, *t*(179) = −1.85, *p* = 0.067, *n* = 184 axons) or between the Open and Occluded sides of Naris Occlusion animals (*t*-test, *t*(211) = −0.591, *p* = 0.555, *n* = 305 axons). The main groups (Control and Naris Occlusion) were also not significantly different from one another (*t*-test, *t*(388) = −1.32, *p* = 0.19, *n* = 489 axons) (***Figure 5***, row C).

Together, we found no evidence of changes in oligodendrocyte lineage cells or myelinated axons in response to 30 days of UNO in adult animals, despite the system wide changes in nodes of Ranvier. This indicates that nodes of Ranvier may change independently of the differentiation and myelination of new OPCs or pre-existing myelin (see Discussion).

### Mitral cell firing properties change following unilateral naris occlusion

Sensory deprivation and enrichment can have dramatic effects on axon morphology and neuronal firing properties (***Jamann et al., 2021***; ***Kuba et al., 2010***; ***Evans et al., 2015***). We have described subtle length changes in the AIS and nodes of Ranvier following UNO, but do these changes affect MC physiology?

To assess MC physiology, we performed whole cell current clamp recordings on UNO and Control mice. Cells were held at −60mV for all experiments. Membrane resting potential was not significantly different between the Control and Naris Occlusion groups (Control −52.9 ± 0.26mV, *n* = 23 cells, Naris Occlusion −52.5 ± 0.423mV, *n* = 19 cells, *t*-test, *t*(32.3) = −0.155, *p* = 0.878, presented as mean ± standard error of the mean [SEM]), or between the Control, Open, and Occluded groups (Control −52.9 ± 0.256mV, *n* = 23 cells, Occluded = −52.8 ± 0.76mV, *n* = 12 cells, Open = −52 ± 0.915mV, *n* = 7 cells, ANOVA, *F*(2,39) = 0.0436, *p* = 0.957, presented as mean ± SEM). MCs are involved in complex circuits and receive both excitatory and inhibitory inputs which affects firing patterns and synchrony (***Schoppa and Westbrook, 2001***; ***Egger and Urban, 2006***; ***Fukunaga et al., 2014***). To isolate MCs from OB circuits and measure intrinsic firing patterns, we performed all recordings in the presence of the glutamatergic inhibitors 6,7-dinitroquinoxaline-2,3-dione (DNQX, 10μM), 2-amino-5-phosphonopentanoic acid (APV, 50μM), and the GABAergic inhibitor gabazine (5μM). MCs have large, complex dendrites and axons which can easily be damaged during the slicing procedure. To ensure damaged MCs were not influencing recordings, we performed a subset of recordings with biocytin in the patch pipette (~2mg/ml), then fixed and immunolabeled the slices for AISs after recording (***Figure 6***A, A’; see Methods). Our data indicate that the majority of recorded cells contained a discernible, AnkG+ AIS and axon (89%, ***Figure 6**A*, white arrow), and a full primary dendrite innervating a glomerulus (72%, ***Figure 6***A, yellow arrow, *n* = 18 cells, 5 Control, 13 Naris Occlusion).

**Figure 6.**
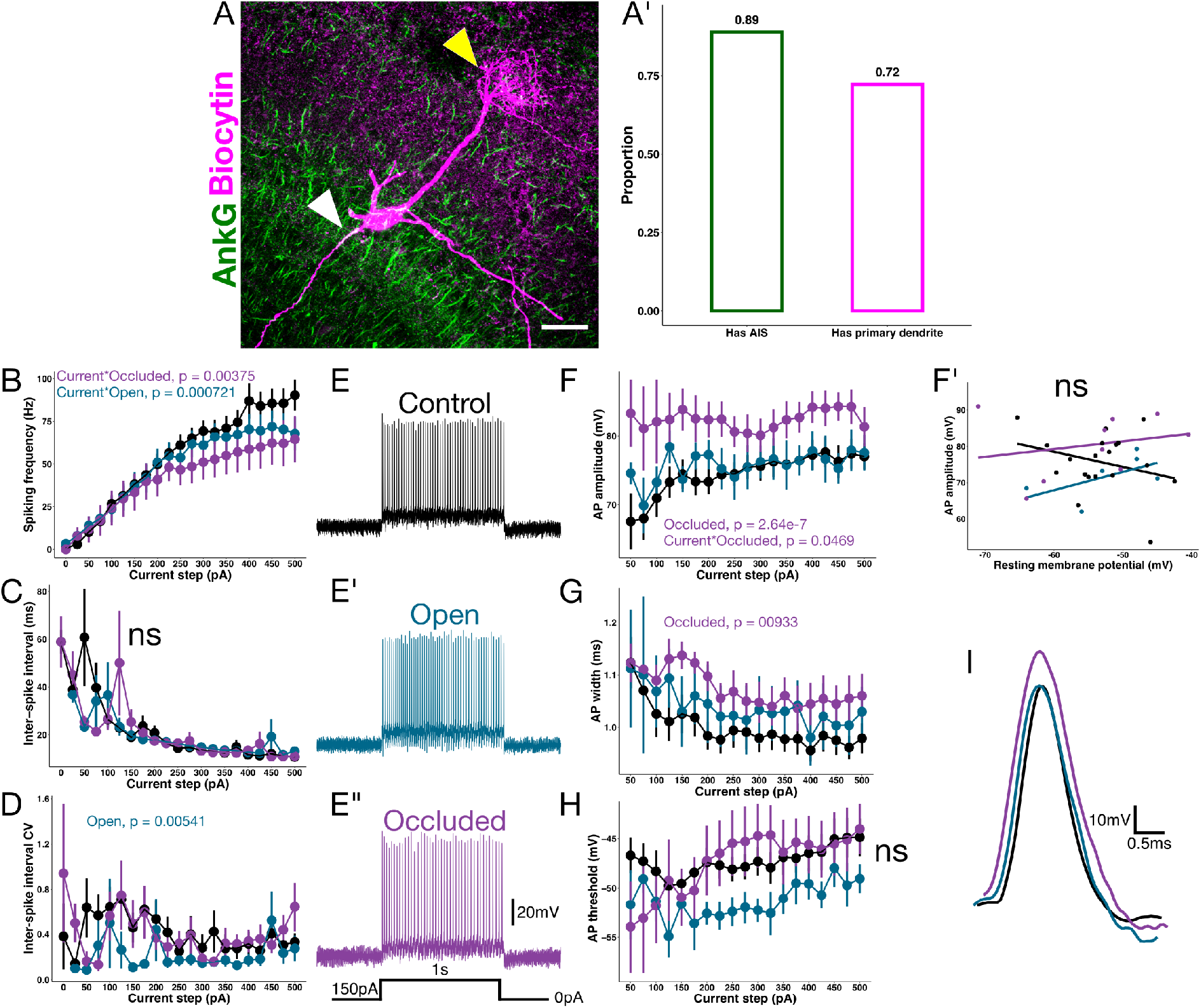
MC spiking and Action potentials (APs) adapt to UNO. (**A**) Example of a MC filled with biocytin (magenta) and immunolabeled for the AIS (AnkG, green). White arrowhead indicates the cell’s AIS, and the yellow arrowhead indicates the primary dendrite innervating a glomerulus. Scale bar is 25μm. (A’) Quantification of structural features of a subset of MCs that were filled and immunolabeled *(n* = 18 cells, 5 Control, 13 Naris Occlusion). 89% of labeled cells had an identifiable AIS, and 72% of cells had an identifiable primary dendrite, indicating that damaged cells likely did not skew results. (**B**) MC spiking frequency across 25pA current steps from 0-500pA. There was a significant interaction between current and Occluded (GLM, Current*Occluded *t* = −2.91, *p* = 0.00375), and current and Open cells vs. Control (GLM, Current*Open, *t* = −3.39, *p* = 0.000721). *P* values for the significant interactions are shown on the plot. (**C**) Inter-spike interval time for current steps, similar to (**B**). The main model was not significantly different from the current-only model, indicating Group had no significant effect on the overall inter-spike interval. (**D**) Plot of coefficient of variation (CV) for inter-spike interval. The Open group had significantly lower CV than control (*t* = −2.79, *p* = 0.00541). *P* value for the Open term is shown on the plot. (**E**) Example current clamp trace from a Control MC in response to a 150pA 1s current injection. (**E’**) Example current clamp trace from an Open MC in response to a 150pA 1s current injection. (**E”**) Example current clamp trace from an Occluded MC in response to a 150pA 1s current injection. (**F**) Plot of AP amplitude across current steps. APs from Occluded MCs had significantly larger amplitudes compared to Control (*t* = 4.27, *p* = 2.23e-5). The Current*Occluded interaction also had a significant effect on AP amplitude (*t* = −1.99, *p* = 0.0469). *P* values for the significant terms are shown on the plot. (F’) Plot of AP amplitude and resting membrane potential at a 150pA step for a subset of cells. There was no significant correlation between amplitude and resting membrane potential for any of the groups in the membrane potential range observed (Pearson’s product-moment correlation, Control, *t*(17) = −1.22, *p* = 0.238, *n* = 18 cells, Open *t*(4) = 1.39, *p* = 0.236, *n* = 5 cells, Occluded *t*(7) = 0.626, *p* = 0.551, *n* = 8 cells). (**G**) Plot of AP width in response to current steps. APs from Occluded bulb MCs were significantly wider than Control (GLM, *t* = 2.61, *p* = 0.00933). *P* value for the Occluded term shown on the plot. (**H**) Plot of AP threshold in response to current steps. Open bulb MCs had only marginally lower thresholds than Control (GLM, *t* = −1.92, *p* = 0.0557), but the effect was not significant at *a* = 0.05. (**I**) Example traces of single action potentials from a Control bulb MC (black), Open bulb MC (blue), and an Occluded bulb MC (magenta) in response to a 150pA current step. *N* cells = 23 Control, 13 Occluded, and 7 Open for all experiments unless otherwise noted. All plots are mean ± standard error of the mean

MCs are diverse, both in terms of morphology and physiology, displaying both bursting and regular spiking patterns in response to current injections (***Padmanabhan and Urban, 2014**,**2010***; ***Fadool et al., 2011***; ***Chen and Shepherd, 1997***). We first investigated the spiking behavior of MCs in response to a series of current injections from 0-500pA with a step size of 25pA (***Figure 6***B). To test whether spiking patterns were different, we fit generalized linear models (GLMs) to the spike frequencies. The main model was of the form:

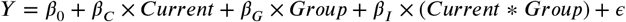

Where *Y* is the dependent variable (in this case the number of spikes), *β*_0_ is the intercept, *β_C_* is the injected *Current, β_G_* is *Group*, and includes Control, Open, or Occluded, *β*_I_ are the *Current*Group* interaction terms between *Current* and *Group*, and *ε* is the error term. We used a Gaussian family link function when fitting the GLMs. To compute the significance of the main model, we used an Analysis of Deviance table to compare it to a GLM containing only current as the independent variable *(Y = β*_0_ + *β_c_* × *Current* + *ε*).

For MC spiking, the main model was significant compared to the current-only model (***Figure 6***B, *F* = 8.57, *p* = 8.7e-7). In the main model, we found that firing rates increased significantly as a function of current (GLM, *t* = 20.5, *p* < 2e-16). Additionally, the interactions between current and Occluded, and current and Open had a significant effect on firing rates vs. Control (GLM, Cur-rent*Occluded interaction *t* = −2.91, *p* = 0.00375, Current*Open interaction *t* = −3.39, *p* = 0.000721, *n* cells = 23 Control, 13 Occluded, and 7 Open; for all GLM results, the reported *t* and *p* values are rounded to 3 significant figures in the text). See Table 1 for the model summary, and supplemental data set 1 for pairwise comparisons between groups.

**Table 1.**
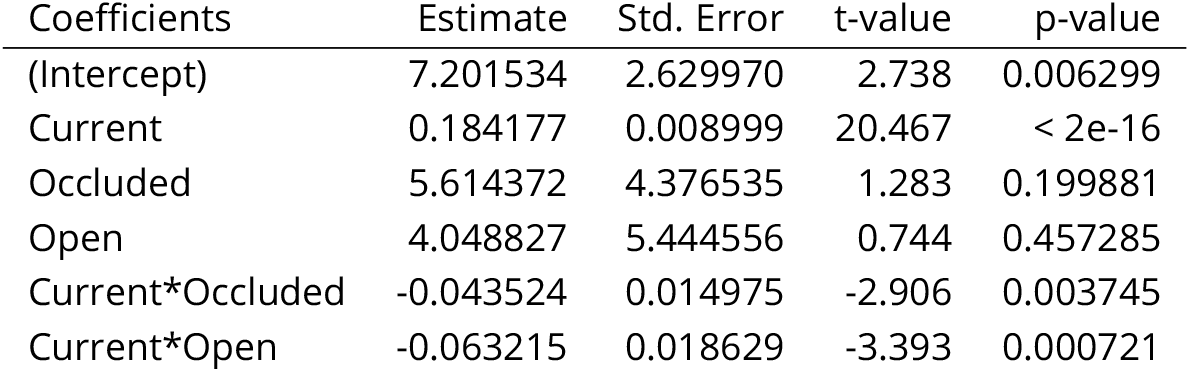
Spike count GLM summary.

Previous computational modeling and experimental studies have indicated that spike timing (measured via inter-spike interval or onset threshold) is sensitive to minor changes in AIS length (***Evans et al., 2015***; ***Baalman et al., 2013***; ***Jamann et al., 2021***). Furthermore, voltage gated potassium channels (K_v_), known to be present at the AIS, strongly influence spiking diversity, AP shape, and AP reliability (***Padmanabhan and Urban, 2014***; ***Kole et al., 2007***; ***Debanne, 2004***). To test whether the MC morphological adaptations were reflected in spiking patterns, we next measured the mean inter-spike interval at each current step. We found that the main model was not significantly different from the current-only model (***Figure 6***C, *F* = 0.848, *p* = 0.495). Only current was significant in the main model, with inter-spike interval decreasing as a function of increasing current (GLM, *t* = −10.5, *p* < 2e-16). There were no significant interactions between any group and current compared to Control (see Table 2 for the model summary, and supplemental data set 2 for pairwise comparisons between groups; *n* cells = 23 Control, 13 Occluded, and 7 Open).

**Table 2.**
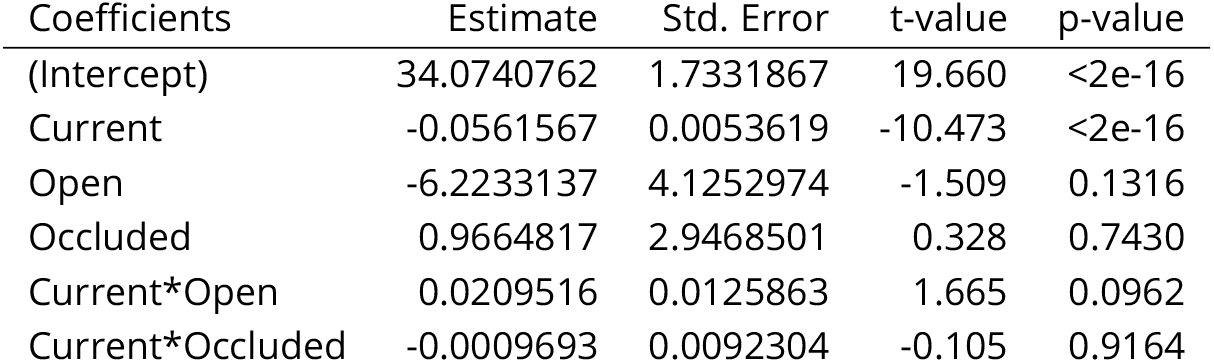
Inter-spike interval GLM summary.

4-Aminopyridine(4AP) sensitive voltage gated potassium channels are implicated in bursting/spiking variability in MCs (***Balu et al., 2004***; ***Padmanabhan and Urban, 2014***). To test whether MCs had altered spiking variability, we calculated the coefficient of variation (CV), a ratio of the SD / mean for inter-spike interval (***Padmanabhan and Urban, 2014**, **2010***). The main model of inter-spike interval CV was significantly different from the current-only model (***Figure 6***D, *F* = 3.27, *p* = 0.0114). Inter-spike interval CV decreased significantly as a function of increasing current (GLM, *t* = −3.28, *p* = 0.0011), and the Open group had significantly lower CV than Control (GLM, *t* = −2.79, *p* = 0.00541; see Table 3 for model summary, and supplemental data set 3 for pairwise comparisons between groups; *n* cells = 23 Control, 13 Occluded, and 7 Open).

**Table 3.** Inter-spike interval coefficient of variation GLM summary.

| Coefficients | Estimate | Std. Error | t-value | p-value |
| --- | --- | --- | --- | --- |
| (Intercept) | 0.6009139 | 0.0665446 | 9.030 | < 2e-16 |
| current | -0.0006841 | 0.0002087 | -3.278 | 0.00110 |
| Open | -0.3988833 | 0.1429577 | -2.790 | 0.00541 |
| Occluded | -0.1390394 | 0.1107063 | -1.256 | 0.20955 |
| Current*Open | 0.0007541 | 0.0004433 | 1.701 | 0.08940 |
| Current*Occluded | 0.0004301 | 0.0003510 | 1.225 | 0.22088 |

We next investigated the kinetics of individual APs in the Control, Open, and Occluded groups. We extracted the first AP from spike trains in current steps from 50pA-500pA (leaving out the lower current steps where few spikes were evoked) to measure AP kinetics. We again fit GLMs to compare relationships.

The main model for AP amplitude (measured from threshold voltage to peak voltage) was significantly different from the current-only model (***Figure 6**F*, *F* = 21.9, *p* < 2.2e-16). AP amplitude increased significantly as a function of increasing current in the main model (GLM, *t* = 4.27, *p* = 2.23e-5). APs from Occluded bulb MCs had significantly larger amplitudes than Controls (GLM, *t* = 5.21, *p* = 2.64e-07). There was also a significant interaction between Current and Occluded vs. Control (GLM, Current*Occluded, *t* = −1.99, *p* = 0.0469). These results are consistent with previous sensory deprivation studies in the chick auditory brainstem (***Kuba et al., 2010***). See Table 4 for the model summary and supplemental data set 4 for pairwise comparisons, *n* cells = 23 Control, 13 Occluded, and 7 Open. To control for different resting potentials possibly affecting AP amplitude, we plotted cell resting potential vs. amplitude for a subset of cells in each group. There was no significant correlation between amplitude and resting membrane potential for any of the groups in the membrane potential range observed (***Figure 6**F*, Pearson’s product-moment correlation, Control *t*(17) = −1.22, *p* = 0.238, *n* = 18 cells, Open *t*(4) = 1.39, *p* = 0.236, *n* = 5 cells, Occluded *t*(7) = 0.626, *p* = 0.551 *n* = 8 cells), which is consistent with previous results (***Balu et al., 2004***).

**Table 4.** AP amplitude GLM summary.

| Coefficients | Estimate | Std. Error | t-value | p-value |
| --- | --- | --- | --- | --- |
| (Intercept) | 70.362952 | 1.222667 | 57.549 | < 2e-16 |
| Current | 0.015724 | 0.003680 | 4.272 | 2.23e-05 |
| Open | 3.690420 | 2.560442 | 1.441 | 0.1500 |
| Occluded | 11.114391 | 2.135462 | 5.205 | 2.64e-07 |
| Current*Open | -0.009227 | 0.007729 | -1.194 | 0.2330 |
| Current*Occluded | -0.012735 | 0.006397 | -1.991 | 0.0469 |

AP width (measured as full width at half maximum) is sensitive to changes in K_v_ channel composition at the AIS, and influences downstream synaptic efficiency (***Kole et al., 2007***; ***Debanne, 2004***). The main model for AP width was significantly different from the current only model (***Figure 6***G, *F* = 11.6, *p* = 4.4e-9). Current was significant in the main model (GLM, *t* = −3.17, *p* = 0.00159), and APs from Occluded MCs were significantly wider than Control (GLM, *t* = 2.61, *p* = 0.00933). There were no significant interactions between group and current (see Table 5 for the model summary, and supplemental data set 4 for pairwise comparisons; *n* cells = 23 Control, 13 Occluded, and 7 Open).

**Table 5.** AP width GLM summary.

| Coefficient | Estimate | Std. Error | t-value | p-value |
| --- | --- | --- | --- | --- |
| (Intercept) | 1.042e-03 | 1.785e-05 | 58.399 | < 2e-16 |
| Current | -1.704e-07 | 5.372e-08 | -3.171 | 0.00159 |
| Open | 3.636e-05 | 3.737e-05 | 0.973 | 0.33099 |
| Occluded | 8.129e-05 | 3.117e-05 | 2.608 | 0.00933 |
| Current*Open | 1.557e-08 | 1.128e-07 | 0.138 | 0.89027 |
| Current*Occluded | -9.298e-09 | 9.338e-08 | -0.100 | 0.92071 |

AP threshold marks the point where an AP becomes an all-or-none response. We defined AP threshold as the voltage when the derivative of the rising phase of the AP reached 25V/s. AP threshold is typically inversely related to AIS length, with longer AISs often displaying lower thresholds (***Kuba et al., 2010***; ***Jamann et al., 2021***). AP threshold is also affected by channel composition ***(Katz et al., 2018***). The main model for AP threshold was significantly different from the current only model (***Figure 6***H, *F* = 6.98, *p* = 1.67e-5). Current had a significant effect on threshold in the main model (GLM, *t* = 2.72, *p* = 0.0067). Surprisingly, APs from Open MCs were only marginally lower than Control (GLM, *t* = −1.92, *p* = 0.0557). There were no significant interactions between group and current (see Table 6 for the model summary, and supplemental data set 4 for pairwise comparisons; *n* cells = 23 Control, 13 Occluded, and 7 Open).

**Table 6.** AP threshold GLM summary.

| Coefficient | Estimate | Std. Error | t value | P value |
| --- | --- | --- | --- | --- |
| (Intercept) | -4.981e+01 | 1.082e+00 | -46.032 | < 2e-16 |
| Current | 8.864e-03 | 3.257e-03 | 2.721 | 0.00668 |
| Open | -4.344 | 2.266 | -1.917 | 0.05571 |
| Occluded | -1.672 | 1.890 | -0.885 | 0.37664 |
| Current*Open | 8.463e-04 | 6.840e-03 | 0.124 | 0.90157 |
| Current*Occluded | 6.891e-03 | 5.661e-03 | 1.217 | 0.22399 |

## Discussion

In the OB, trains of APs at the gamma frequency (oscillations) are generated in response to odors (***Li and Cleland, 2017***; ***Eeckman and Freeman, 1990***; ***Kashiwadani et al., 1999***; ***Bathellier et al., 2006***). These oscillations must reliably travel large distances for further processing in regions such as the piriform cortex, where precise, synchronized arrival determines whether a cell fires (***Nagayama et al., 2010***; ***Franks and Isaacson, 2006***; ***Luna and Schoppa, 2008***; ***Bolding and Franks, 2018***). Despite the importance of reliable AP transmission over large distances, little is known about the myelinated axons that generate and propagate APs in the OB, and whether they adapt in response to changes in sensory input. Here, we characterized the myelinated axons of MCs in the OB and LOT, and tested whether they adapt to altered sensory input using UNO. We found that 30 days of adult UNO led to an increase in AIS length of 8% (~2μm) in MCs from Open bulbs relative to both Occluded bulbs and Control (***Figure 4***). A change of similar magnitude is predicted by computational modeling to result in significant physiological changes in nonlinear AP characteristics (***Baalman et al., 2013***). Indeed, we found Naris Occlusion had a significant effect on spiking patterns, reducing spiking frequency and spike variability (***Figure 6***B, D; Tables 1 and 3).

In addition to overall spiking patterns, whole cell patch clamp also revealed significant differences in AP width in AISs from the Occluded side, despite the Occluded cells AIS length being similar to Controls (***Figure 3*** and ***Figure 6***). Why would AP width increase in Occluded MCs? One possibility is an adaptation in K_v_ channel composition or number. K_v_ channels present at the AIS are known to regulate AP width (***Kole et al., 2007***; ***Debanne, 2004***), and have significant effects on MC firing pattern diversity (***Padmanabhan and Urban, 2014**, **2010***). More K_v_ channels could lead to wider action potentials, which increase the probability of depolarization of downstream cells (***Kole et al., 2007***). This may translate to a counterintuitive *increase* in odor sensitivity on the Occluded side due to a higher probability of depolarizing a downstream cell in the piriform cortex. This hypothesis is supported by a previous study showing that Naris Occlusion mice outperformed Control mice in a habituation-dishabituation olfactory discrimination task (***Angely and Coppola, 2010***). Our results could represent a novel mechanism to explain that finding. Future work will determine whether AP shape changes measured in Occluded bulbs translate to physiological adaptations in synapses at the piriform cortex.

Another surprising finding was the different distributions of AIS lengths. AIS length distributions were significantly different between Control animals and Naris Occlusion animals. This increased diversity was counter-intuitively reflected in an overall lower inter-spike interval CV in the Open group compared to Control (***Figure 6***D, Table 3). What explains this increased length variance and decreased spiking variance? MCs are biophysically diverse cells, with cell-level firing differences largely attributed to local circuits and OB oscillations, as well as ion channel diversity, particularly in K_v_ channels (***Padmanabhan and Urban, 2010**,**2014***; ***Heyward et al., 2001***; ***Angelo and Margrie, 2011***). Spiking diversity is thought to increase the information carrying capacity of neurons by de-correlating the firing patterns of groups of neurons (***Padmanabhan and Urban, 2010***). One hypothesis is that the increased range of AIS lengths in Naris Occlusion animals reflects an attempt to encode more information with half the resources. The less variable spiking patterns we measured *in vitro* are likely different than MC spiking patterns *in vivo. In vivo*, respiratory rhythm, local circuits/oscillations, and intrinsic membrane oscillations lead to different MC spiking patterns than *in vitro (**Cang and Isaacson, 2003***; ***Li et al., 2017***; ***Cury and Uchida, 2010***; ***Angelo and Margrie, 2011***). Further experiments will determine how neuronal spike rates change over the course of UNO *in vivo.*

AISs in the Open bulbs are significantly longer than AISs in Control or Occluded bulbs (***Figure 4***D). Longer AISs are typically associated with lower firing thresholds (***Kuba et al., 2010***; ***Jamann et al., 2021***). We found that threshold was only marginally lower across current steps in Open bulb MCs (***Figure 6***, *p* = 0.0557, see Table 6). While not significant at *α* = 0.05, increased AIS length and marginally decreased firing threshold on the Open side (the side still receiving input) was somewhat surprising. Previous studies report that neurons undergoing deprivation had longer AISs and lower firing thresholds compared to Controls (***Kuba et al., 2010***; ***Jamann et al., 2021***). Similarly, neurons chronically stimulated or in an enriched environment respond by decreasing AIS length or translocating AISs away from the soma, with the effect of decreasing excitability ***(Evans et al., 2015; Grubb and Burrone, 2010***; ***Evans et al., 2013***; ***Jamann et al., 2021***). These changes have typically been interpreted in light of a homeostatic response by neurons to regulate excitability (***Wefelmeyer et al., 2016***; ***Yamada and Kuba, 2016***). The opposite effects in our study may be more analogous to a synaptic Hebbian-like response (***Abbott and Nelson, 2000***) to increased input, where neurons become more excitable because they must now work double duty to encode the same amount of information. Previous work in inhibitory dopaminergic neurons in the OB has also described this “inverted” plasticity of the AIS in response to chronic stimulation, noting a cell-type specific response (***Chand et al., 2015***).

Nodes of Ranvier allow for fast saltatory conduction along myelinated axons by regenerating propagating APs at successive gaps in the myelin sheath (***Susuki et al., 2013***; ***Castelfranco and Hartline, 2016***). Recent work has implicated nodes of Ranvier as potential sites for plasticity ***(Ford et al., 2015; Arancibia-Cárcamo et al., 2017***; ***Dutta et al., 2018***; ***Cullen et al., 2021***). We measured nodes of Ranvier in the LOT following 30 days of UNO. In contrast to the AIS, we found no significant differences in mean node of Ranvier length within Control or within Naris Occlusion animals ***(Figure 4***). Surprisingly, we found that node lengths were significantly different when comparing Naris Occlusion animals (both Open and Occluded sides) to Controls (both Left and Right sides). While it is unclear why Naris Occlusion animals exhibit different node lengths compared to Controls, bilateral adaptations following UNO are not unprecedented. In particular, a study investigating *in vivo* synaptic responses of olfactory receptor neurons following UNO found that odor-induced synaptic release was equally reduced in the Open and Occluded sides compared to Controls ***(Kass et al., 2013***). Similarly, gene expression in the olfactory mucosa changes on both the Open and Occluded sides compared to Controls (***Coppola and Waggener, 2012***). Bilateral changes following UNO highlight the importance of using comparisons with separate Control animals in UNO studies in addition to within animal (e.g. Left vs. Right bulb) comparisons (***Coppola, 2012***).

Despite finding no changes in the myelin sheath (measured by *g*-ratio) or oligodendrocyte lineage cells (***Figure 5***), we found that nodes in Naris Occlusion animals were significantly shorter than nodes from Control animals (***Figure 4***). In the central nervous system, nodes of Ranvier form via a complex interaction between the myelin sheath (made by oligodendrocytes) and neuronal axons (***Susuki et al., 2013***), so the lack of change in oligodendrocyte lineage cells and myelin was surprising. Previous work in the visual system reported monocular deprivation caused proliferation and maturation of oligodendrocytes, as well as a higher number of short myelin sheaths on the deprived side (***Etxeberria et al., 2016***). Motor learning, somatosensory stimulation, and optogenetic stimulation of neuronal activity in motor cortex have also been reported to increase oligodendrocyte lineage cell proliferation, differentiation, and cause myelin sheath remodeling ***(Gibson et al., 2014; McKenzie et al., 2014***; ***Xiao et al., 2016***; ***Hughes et al., 2018***; ***Hill et al., 2018***). Many cases of reported myelin plasticity have involved *in vivo* studies limited to the upper two layers of cortex, a region known to be more sparsely myelinated than large axonal tracts like the LOT ***(Tomassy et al., 2014***). In studies where changes in myelination and oligodendrocytes occurred in adulthood, they were most dramatic on axons which were intermittently myelinated (***Hill et al., 2018***; ***Hughes et al., 2018***), or in animals undergoing active learning (***McKenzie et al., 2014***; ***Xiao et al., 2016***). It is possible that more highly myelinated regions like LOT may be more stable, and changes in oligodendrocyte lineage cell proliferation and differentiation may require active learning rather than just reduced input.

How could shorter nodes of Ranvier affect olfaction? Modeling studies indicate that shorter nodes are associated with slower conduction velocity (***Arancibia-Cárcamo et al., 2017***), and a recent study found that competency at a spatial learning task was associated with longer nodes of Ranvier in the hippocampus and faster conduction speed compared to controls with shorter nodes (***Cullen et al., 2021***). Based on these results, one would assume that conduction velocity is slower in animals that underwent Naris Occlusion compared to Controls. It is not immediately clear what effect potentially slower conduction velocity would have on olfaction. While speed is not thought of as essential to olfactory processing, coordinated signals and the reliable transmission of oscillations are associated with olfactory processing and learning (***Franks and Isaacson, 2006***; ***Laurent et al., 1996; Kay et al., 2009***; ***Losacco et al., 2020***). Since the node of Ranvier adaptations appear global (differing between Control and Naris Occlusion but not between LOT sides), one possibility is that slowing of AP transmission between the OB and downstream olfactory processing regions serves to better correlate disparate olfactory signals, allowing a higher probability of depolarizing downstream cells. However, it is hard to predict how adaptations in myelinated axons will affect the system-level information transfer in oscillations. Computational modeling studies emphasize the importance of myelin and conduction speed tuning for oscillation synchrony ***(Pajevic et al., 2014***), and our lab has previously shown that mild myelin disruption in the *Plp1-null* mouse leads to increased oscillatory power in the theta and beta frequencies (***Gould et al., 2018***). Future work will elucidate the effects of smaller nodes in Naris Occlusion animals on conduction velocity and downstream signal integration. Ourwork provides evidence fora novel form of cellular plasticity in the olfactory system where myelinated axons adapt to changing experience input in adult animals. It is unclearwhatthe systems level consequences of these novel adaptations are, but they are likely to have important consequences for downstream olfactory system information processing.

## Methods

### Source code and imaging data

Source code used for statistics, figure generation, and analysis can be found in the following repositories:

- https://github.com/nkicg6/excitable-axonal-domains-figures
- https://github.com/nkicg6/excitable-axonal-domains-physiology
- https://github.com/Macklin-Lab/imagej-microscopy-scripts

Source imaging data can be found in the following OSF project:

- https://osf.io/ez3qt/

### Statistics

All statistics were performed using the R programming language (***R Core Team, 2019***), and several packages from the tidyverse family (***Wickham et al., 2019a***). Plots were made using the R package ggplot2 (***Wickham, 2016***), using Cairo (***Urbanek and Horner, 2020***) for PDF export, or the Python package matplotlib (***Hunter, 2007***). The R packages dplyr (***Wickham et al., 2019b***), readr ***(Wickham et al., 2018***), tidyr (***Wickham and Henry, 2019***), and cowplot (***Wilke, 2019***) were used for data processing and analysis. Results are given as mean ± SD, unless otherwise noted in the text. Values were compared with the Welch two sample *t*-test using R. In the case of multiple comparisons, p-values were corrected using FDR (***Benjamini and Hochberg, 1995***). Statistical significance was set at *a* = 0.05 (*p* ≤ 0.05).

The results of statistical tests are presented with the test statistic, degrees of freedom, and p value, if applicable. For example, a *t*-test is presented as: *t*-test, *t*(degrees of freedom) = t-statistic, *p* = p-value. Test statistics and *p*-values are rounded to 3 significant figures in the text.

### Animal care

Adult P60-90 wild-type male and female mice (strain C57BL/6J, Jackson Lab #000664) were used for all experiments. Mice were bred and housed in the University of Colorado Anschutz Medical campus vivarium.

Animals were always housed in single-sex cages of 2-5 individuals with a 14/10 hour light/dark cycle. Mouse chow and water were available ad libitum. Experimental protocols and animal care were performed in accordance with the Institutional Animal Care and Use Committee at the University of Colorado Anschutz Medical Campus.

### Unilateral naris occlusion

Adult (~P60) wild-type mice (males and females) were anesthetized with an intraperitoneal injection of 100mg/kg ketamine, 10mg/kg of xylazine. When unresponsive to a toe pinch, animals were given a local application of 2% lidocaine to the external right naris, and the right naris was briefly cauterized with a Bovie high temperature cautery (Bovie Medical Corporation, Clearwater, Florida) and a small amount of super glue was applied to seal the naris. We applied gentamycin ophthalmic ointment to the eyes during surgery to maintain hydration. After the surgery, animals were given 0.4ml of sterile saline subcutaneously (SC), and SC carprofen (10mg/kg) the day of surgery and the day after. We monitored the animals as they recovered from anesthesia on a heating pad before returning them to the vivarium in accordance with the University of Colorado Anschutz Medical Campus Institutional Animal Care and Use Committee.

Before the animals were sacrificed for immunohistochemistry, EM, or physiology, we confirmed naris occlusion by placing 0.1% Triton X-100 on the Occluded naris and ensuring that no bubbles formed (bubbles would indicate incomplete occlusion). A subset of animals were fixed and stained for TH to confirm effective UNO ***Figure 3***.

Control animals (also called sham) were cage mates of Occluded animals and underwent the exact same protocol sans cauterization and closing the naris.

### Immunohistochemistry sample preparation

Mice were anesthetized with Fatal-Plus (Vortech Pharmaceuticals, Dearborn, Michigan) and tran-scardially perfused with 20ml of 0.01M phosphate buffered saline (1X PBS) followed by 20ml of 4% paraformaldehyde (diluted from a 32% aqueous paraformaldehyde solution with PBS; Electron Microscopy Sciences [EMS], Hatfield, Pennsylvania) at a flow rate of 10-14ml/min. The brains were carefully removed and post-fixed in 4% paraformaldehyde for 1-2 hours at 4°C. Following post-fix, brains were placed in a 30% sucrose-PBS solution for 48 hours for cryoprotection. Brains were then embedded in molds in optimal cutting temperature (OCT; EMS, Hatfield, Pennsylvania) and frozen at −80°C until sectioning.

Slices were serially sectioned in the horizontal plane at 30-40μm thick using a Leica CM1950 (Leica Biosystems, Buffalo Grove, Illinois) and collected as free floating sections in PBS in 24 well plates. We performed immunohistochemistry within 7-14 days of sectioning.

### Immunohistochemistry

#### Section sampling

We followed the principles of unbiased stereology for cell number and AIS/node of Ranvier length quantification (***Mouton, 2002***; ***Mouton et al., 2017***). We defined the anatomical quantification area to be the appropriate regions of OB and LOT (MCL for AISs, GCL and LOT for oligodendrocyte lineage cell quantification, and GL for TH quantification) between approximately-2.04mm to −5.64mm ventral to bregma suture (***Franklin and Paxinos, 2013***; ***Capra, 2003***). We used a systematic random sampling scheme to ensure unbiased cell number and AIS/node of Raniver length quantification (***Mouton, 2002***). Briefly, we counted the number of sections collected for each animal in the target anatomical region, and used the Python programming language (Python 3.7.0) “random.choice” function to choose a random starting point for each set. We then chose every *n*^th^ section, where *n* = the number of slices in the anatomical area divided by the target sampling number of slices (typically 4-5 sections/animal).

#### Section labeling

We performed immunohistochemical labeling using a free-floating slices protocol modified from previous studies (***Gould et al., 2018***; ***Ahrendsen et al., 2018***). Slices were processed in batches to reduce variability and facilitate comparison.

Slices were washed 3 times for 5 minutes each in 1X PBS, then placed in 10mM sodium citrate 0.05%Tween-20 buffer (pH 6) for 1 minute to equilibrate before microwave-based antigen retrieval in a PELCO BioWave Pro microwave (550W for 5 minutes; Ted Pella, Redding, California). Following antigen retrieval, slices were washed 3 times for 5 minutes each in 1X PBS, and permeabilized slices with 0.1-0.3% Triton X-100 PBS solution for 20 minutes. Slices were then blocked the slices for 1 hour in a solution of 5% normal goat or donkey serum (depending on the antibodies) in 0.3% Triton X-100 PBS at room temperature. Slices then incubated overnight on a rocker at room temperature in the blocking solution + primary antibodies (see Table 7 for antibody concentration information, and Table 9 for vendor information). Incompatible antibodies (e.g., TH and Caspr) were not used on the same slices.

**Table 7.** Primary antibody information.

| Antigen | Host/isotype | Concentration (μl antibody : μl block) |
| --- | --- | --- |
| AnkG | Mouse/IgG2a | 1:500 |
| Plp | Rat | 1:1000 |
| TH | Mouse/IgG1 | 1:500 |
| Caspr | Mouse/IgG1 | 1:500 |
| Na<sub>v</sub>1.6 | Rabbit | 1:500 |
| Olig2 | Mouse/IgG2a | 1:500 |
| PDGFRα | Rat | 1:500 |
| CC1 | Mouse/IgG2b | 1:500 |

The following day, slices were washed 4 times for 10 minutes each in 1X PBS, then incubated for 2 hours in blocking solution and secondary antibodies on a rocker at room temperature (see Table 8 for secondary antibody concentrations, and Table 9 for vendor information). Following secondary antibody incubation, slices were washed 4 times for 5 minutes in 1X PBS, the incubated with the nuclear label Hoechst (diluted 1:5000 in 1X PBS) for 2 minutes on a rocker at room temperature. If slices were labeled with red fluorescent Nissl (NeuroTrace, Thermo Fisher, Waltham, Massachusetts), we incubated them in a PBS-Nissl solution in the dark for 1 hour, then washed 4 times for 5 minutes in 1X PBS before performing the Hoechst label.

**Table 8.** Secondary antibody information.

| Antigen | Secondary antibody | Concentration (μl antibody : μl block) |
| --- | --- | --- |
| AnkG | Goat anti IgG2a | 1:1000 |
| Plp | Goat anti Rat | 1:1000 |
| TH | Goat anti IgG1 | 1:1000 |
| Caspr | Goat anti IgG1 | 1:1000 |
| Na<sub>v</sub>1.6 | Goat anti Rabbit | 1:1000 |
| Olig2 | Goat anti IgG2a | 1:1000 |
| PDGFRα | Goat anti Rat | 1:1000 |
| CC1 | Goat anti IgG2b | 1:1000 |
| Hoechst | NA | 1:5000 |
| NeuroTrace (Nissl) | NA | 1:250 |

Slices were then washed 2 times for 5 minutes each in 1X PBS and transferred to 0.01M phosphate buffer for mounting. Slices were mounted on uncharged Gold Seal Rite-on glass slides (Thermo Fisher, Waltham, Massachusetts, CAT# 3050) using Fluoromount-G mounting media (SouthernBiotech, Birmingham, Alabama) and #1.5 coverslips (Thermo Fisher, Waltham, Massachusetts). Slides were then stored in the dark at 4°C until imaging.

#### Physiology section labeling

Following whole cell patch clamp experiments, a subset of sections were fixed in 4% paraformaldehyde for 2 hours, washed in 1X PBS, and labeled for biocytin, AnkG, and nuclei. After fixation, slices were washed 3 times for 5 minutes in 1X PBS, then permeabilized in 0.3% Triton X-100 PBS for 20 minutes. Slices were then blocked for 1 hour in a solution of 5% normal goat serum in 0.3% Triton X-100 PBS. Primary antibodies against AnkG were diluted in blocking solution (see Table 7), and slices were left at 4°C on a rocker for approximately 72 hours. Next, slices were washed 4 times for 10 minutes in 1X PBS, before secondary antibodies against AnkG (see Table 8) and Alexa Fluor 594-conjugated Streptavidin (1 μl Streptavidin to 500 μl block) were diluted in blocking solution and added to the slices. Slices incubated in the dark in secondary antibody at 4°C for 48 hours. Following secondary antibody incubation, slices were washed 2 times for 5 minutes in 1X PBS, and incubated in Hoechst diluted in PBS (see Table 8) for 5 minutes. Next, slices were washed 3 times for 30 minutes in 1X PBS, and mounted on uncharged Gold Seal Rite-on glass slides (Thermo Fisher, Waltham, Massachusetts, CAT # 3050) using ProLong Gold Antifade mounting media (Thermo Fisher, Waltham, Massachusetts, CAT # P10144) and #1.5 coverslips (Thermo Fisher Scientific, Waltham, Massachusetts). Slices were cured in ProLong Gold on a flat surface for 48 hours in the dark before imaging.

### Electron microscopy fixation and sample preparation

Mice were anesthetized with Fatal-Plus (Vortech Pharmaceuticals, Dearborn, Michigan) and transcardially perfused with 10ml of 1X PBS followed by 30ml of EM fixative (2.5% paraformaldehyde, 2.5% glutaraldehyde, 2mM calcium chloride, 0.1M sodium cacodylate buffer). The PBS and EM fixative were kept on ice at 4°C. After perfusion, the brain was removed and post fixed for ~12 hours at 4°C in the EM fixative. Following post-fix, we stored the brains in 0.1M sodium cacodylate buffer at 4°C until sectioning. For sectioning, brains were embedded in 4% low-melt agarose for stability and cut in 1X PBS on a vibratome (Ted Pella, Redding, California) into 300μm thick coronal sections encompassing the region 2.45-3.05mm anterior to the bregma suture. Further processing was performed as previously described (***Ahrendsen et al., 2018***). Briefly, using a PELCO Biowave Pro Tissue Processor (Ted Pella, Redding, California), the tissue was rinsed in 100 mM cacodylate buffer and then postfixed in a reduced osmium mixture consisting of 1% osmium tetroxide and 1.5% potassium ferrocyanide followed by 1% osmium tetroxide alone. Dehydration was performed in a graded series of acetone dilutions (50%, 70%, 90%, and 100%) containing 2% uranyl acetate for en bloc staining. Finally, tissue was infiltrated and embedded in Embed 812 (EMS, Hatfield, Pennsylvania) and cured for 48 h at 60°C. Tissue was oriented so that sections could be cut in the coronal plane to visualize the LOT. Ultrathin sections (65nm) were mounted on copper slot grids and viewed at 80kV on a Tecnai G2 transmission electron microscope (FEI, Hillsboro, Oregon). Electron micrographs were obtained in consistent regions in the lateral portion of LOT.

### Confocal and Electron Microscopy imaging and quantification

All image analysis (EM and confocal) was performed using the freely available Fiji distribution of ImageJ (***Schindelin et al., 2012***). EM and confocal images were blinded using a custom Fiji script called blind-files, provided as part of the Lab-utility-plugins update site (see Source code and imaging data).

#### AIS quantification

For AIS quantification, images were taken on a Nikon A1R resonance scanning confocal microscope (Nikon, Melville, New York) with a Nikon Plan Fluor 40x oil immersion objective (numerical aperture = 1.3). We acquired 3D confocal stacks 318μm × 318μm × 15μm (X × Y × Z)with a voxel size of 0.31 × 0.31 × 0.225μm^3^ (X × Y × Z). We acquired Hoechst (405nm excitation), AnkG (488nm excitation), Nissl (561nm excitation), and Plp (640nm excitation) images with a line average of 4. We took images from lateral and medial MCL of each bulb for analysis. Images were blinded and analyzed in 3D using the semi-automated tracing tool SNT (***Arshadi et al., 2020***), a Fiji plugin. We traced from the origin of the AnkG signal at the base of the Nissl+ soma to the termination of the AnkG signal (typically ending abruptly in a Plp+ myelin sheath, see ***Figure 1***). We then used the Fit Paths option in

SNT to automatically optimize path fits using 3D intensity around each traced node ***(Arshadi et al., 2020***) before exporting length measurements as comma separated value (CSV) spreadsheets for analysis using R. AIS length analysis was performed for treatment group (Control vs. Naris Occlusion), within group (Left vs. Right for Control, Open vs. Occluded for Naris Occlusion), and between group (Control vs. Open vs. Occluded). When AISs were grouped by animal, we calculated the mean AIS length per section and animal.

#### Tyrosine hydroxylase quantification

Horizontal sections (encompassing both olfactory bulbs) were labeled for TH and nuclei (Hoescht) as described above. We took tiled images of the whole sections (both bulbs) using a Zeiss Axio Imager.M2 widefield microscope with a Zeiss 20x Plan-Apochromat (numerical aperture = 0.8) objective and an HXP 120 metal halide lamp (Carl Zeiss Microscopy, White Plains, New York). The XY pixel size was 0.65μm/pixel. We then cropped the tiled images in Fiji so only one bulb was present in each image and labeled them appropriately. Images were then blinded for analysis. We used the polygon tool in Fiji to trace the GL (using the Hoechst labeled nuclei of glomeruli as a guide), then used a custom Fiji script to extract fluorescence intensity measurements into CSV format for further analysis using R. We calculated a mean fluorescence intensity (sum pixel intensity / area traced) per side and animal for the analysis. Intensity is presented in arbitrary units (a.u.) and either compared directly within animal, or as a relative intensity between Control and Naris Occlusion (Right/Left or Occluded/Open, see ***Figure 3***).

#### Node of Ranvier quantification

Node of Ranvier images were taken on a Nikon A1R resonance scanning confocal microscope (Nikon, Melville, New York) with a Nikon Plan Fluor 40x oil immersion objective (numerical aperture = 1.3) using a 3X optical zoom. We acquired 3D confocal stacks 106μm × 106μm × 15μm (X × Y × Z) with a voxel size of 0.106 × 0.106 × 0.25 μm^3^ (X × Y × Z). We acquired Caspr (488nm excitation), and Na_v_1.6 (561nm excitation) and Plp (640nm excitation) with a line average of 4. We took images from the LOT of both bulbs on each slice for analysis. Images were blinded before we manually traced nodes. We traced a random subset of 10-25 nodes per image in 3D using the Fiji segmented line tool. We only traced nodes where the Na_v_1.6 signal was approximately contained in a single optical section to control for out of plane errors. After tracing, the ROI files of all traced nodes were saved, and we used a custom script to extract the Na_v_1.6 fluorescence signal and fit a Gaussian to the signal using Fiji’s curve fitting tools. We extracted the fit parameters and calculated the full width at half maximum of the fit Gaussian to determine node length. To control for poor fitting, we excluded all nodes whose Gaussian fit R^2^ value were < 0.9. When node lengths were grouped by animal, we calculated the mean node length per section and animal.

#### Oligodendrocyte lineage cell quantification

Oligodendrocyte lineage cell images were taken on a Nikon A1R resonance scanning confocal microscope (Nikon, Melville, New York) with a Nikon Plan Fluor 40xoil immersion objective (numerical aperture = 1.3). We acquired 3D confocal stacks 318μm × 318μm × 10μm (X × Y × Z) with a voxel size of 0.31 × 0.31 × 0.25μm^3^ (X × Y × Z). We acquired Hoechst (405nm excitation), Olig2 (488nm excitation), PDGFR*α* (561nm excitation), and CC1 (640nm excitation) with a line average of 4. We took images from the GCL and LOT of each bulb for analysis. Images were blinded and analyzed in 3D using the Fiji Cell Counter plugin. Only cells completely contained within the imaging field were counted. Cell Counter plugin results were saved as CSV for analysis in R.

#### Whole cell patch clamp cell morphology quantification

We took images of a subset of cells filled with biocytin and stained for AnkG on a Nikon A1R resonance scanning confocal microscope (Nikon, Melville, New York) with a Nikon Plan Fluor 40x oil immersion objective (numerical aperture = 1.3). Some images were tiled to create a representative image of the axon and primary dendrite. Voxel size was 0.62 × 0.62 × 1.1 μm^3^ (X × Y × Z), which was sufficient for a faithful representation of the cells. Images were maximum intensity projected in Z for quantification. We manually checked whether a filled cell had a primary dendrite extending to the glomerulus, and whether it had a visible axon with an AnkG+ AIS.

#### Electron microscopy quantification

EM images were blinded and manually analyzed using the polygon tool in Fiji. We traced a random selection of 10-15 axons per image. The vast majority of axons in the LOT are myelinated by P30 (***Collins et al., 2018***), so we only traced myelinated axons. We traced the axon and myelin sheaths using the polygon selection tool in Fiji, and calculated the *g*-ratio by dividing the perimeter of the axon by the perimeter of the myelin sheath (***Figure 2***). A *g*-ratio of 1 indicates an unmyelinated axon, while computational modeling studies propose an optimal *g*-ratio of ~0.77, balancing energy demands of the structure, space, and axonal conduction (***Chomiak and Hu, 2009***). To calculate axon diameter, we fit the measured perimeter of the polygon to the equivalent circle (perimeter = circumference) and calculated the resulting diameter (Diameter = circumference / *π) **(Ahrendsen et al., 2018***).

### Electrophysiology

#### Acute slice preparation

For optimal patch clamp recordings on older animals (~P90), we used different modified artificial cerebral spinal fluid (ACSF) solutions for slice preparation and slice incubation ***(Ting et al., 2014,2018***). Animals were anesthetized with ketamine/xylazine (100mg/kg ketamine, 10mg/kg xylazine) and when unresponsive they were transcardially perfused with 25ml of ice cold N-methyl-D-glucamine (NMDG) based ACSF (NMDG-ACSF, in mM: 92 NMDG, 2.5 KCl, 1.2 NaH_2_PO_4_, 30 NaHCO_3_ 25 glucose, 20 HEPES, 5 Na-ascorbate, 2 thiourea, 3 Na-pyruvate, 10 MgSO_4_, 0.5 CaCl_2_, adjusted to pH 7.4 with 5M HCl, osmolarity 300-310mmol/kg) bubbled continuously with carbogen (95% oxygen, 5% carbon dioxide). The brain was removed, embedded in 2.5% low melt agarose (diluted in NMDG-ACSF), and cut in 300-400μm horizontal sections using a Compresstome VF-310 (Precision-ary Instruments, Natick, Massachusetts) in carbogen bubbled NMDG-ACSF. Once cut, slices were transferred to incubate at 32°C in carbogen bubbled NMDG-ACSF for 30 minutes (resting period). During the initial resting incubation period, we performed the sodium spike protocol for 3-6 month old mice to optimize gigaohm seal formation (***Ting et al., 2018***). This involved adding set volumes of 2M sodium chloride at regular intervals to slowly re-equilibrate the slices to sodium ions (described in ***Ting et al. (2018)*** Table 2 for 3-6mo mice, in μl: 250 at 5 minutes resting, 500 at 10 minutes resting, 1000 at 15 minutes resting, 2000 at 25 minutes resting, and transfer at 30 minutes).

Following the sodium spike in, slices were transferred to a room temperature HEPES-based ACSF solution for 1 hour before recording (HEPES-ACSF, in mM: 92 NaCl, 2.5 KCl, 1.2 NaH_2_PO_4_, 30 NaHCO_3_, 25 glucose, 5 Na-ascorbate, 2 thiourea, 3 Na-pyruvate, 2 MgSO_4_, 2 CaCl_2_, adjusted to pH 7.4 with 5M NaOH, osmolarity 300mmol/kg). For all solutions, osmolarity was measured with a VAPRO vapor pressure osmometer (Wescor, Logan, Utah).

#### Whole cell patch clamp recording

Whole cell patch clamp was performed with pipettes filled with a potassium gluconate based internal solution (in mM: 130 K-gluconate, 10 HEPES, 10 KCl, 0.1 EGTA, 10 Na2-phosphocreatine, 4 Mg-ATP, 0.3 Na_2_-GTP, adjusted to pH 7.3 with KOH, osmolarity 280mmol/kg). Some recordings were done with 2mg/ml of Biocytin, added the day of the experiment, for post-hoc cell visualization. Pipettes were pulled from borosilicate glass with filaments, inner diameter 0.86mm, outer diameter 1.5mm (item BF-150-86-10, Sutter Instruments, Atlanta, Georgia) to a tip resistance of 3-4MOhms with a P-97 Flaming/Brown type micropipette puller (Sutter Instruments, Atlanta, Georgia).

We performed all recordings in the presence of the glutamatergic inhibitors 6,7-dinitroquinoxaline-2,3-dione (DNQX, 10μM), 2-amino-5-phosphonopentanoic acid (APV, 50μM), and the GABAergic inhibitor gabazine (5μM) in ACSF(in mM: 5 HEPES, 125 NaCl, 2.5 KCl, 1.25 NaH_2_PO_4_, 24 NaHCO_3_,12.5 glucose, 2MgSO_4_, 2 CaCl_2_, adjusted to pH 7.4, osmolarity 300-310mmol/kg, bubbled continuously with carbogen, called recording ACSF).

During recording, slices were placed in a custom perfusion chamber continuously perfused with carbogen bubbled recording ACSF heated to 33-36°C with a SH-27B in line heater and a TC-324C temperature Controller (Warner Instruments, Hollister, Massachusetts). We performed the experiments using a Zeiss Axioskop 2 FS Plus microscope (Carl Zeiss Microscopy, White Plains, New York) equipped with differential interference contrast optics and a 40x (numerical aperture = 0.8) Zeiss Achroplan water immersion objective (Carl Zeiss Microscopy, White Plains, New York). We visualized slices using a CoolSNAP HQ2 camera (Teledyne Photometrics, Tucson, Arizona) with Micro-Manager software version 1.4.22 (***Edelstein et al., 2014***). Patch pipettes were manipulated using a MP-285 manipulator arm driven by a MPC-200 Controller and ROE-200 micromanipulator (Sutter Instruments, Atlanta, GA).

Data were acquired using Clampex software version 10.5.0.9 with an Axopatch 200A amplifier, CV-201A headstage, low pass filtered with a Bessel filter at 2kHz and digitized with an Axon Digidata 1550A at 20kHz (Molecular Devices, San Jose, California). We did not correct forajunction potential. We performed offline filtering of current clamp traces using a 3rd order Savistky-Golay filter with a 0.5ms window. Displayed traces were filtered with a 1ms window for appearance (***Figure 6***).

MCs were identified based on their large cell bodies and position in the MC layer. A subset of cells were filled with biocytin, fixed with 4% paraformaldehyde, and visualized to confirm the presence of an apical dendrite extending to the glomerular layer (***Figure 6***A). Access resistance and resting potential was checked shortly after achieving whole cell configuration and if it exceeded 40MOhms cells were discarded (***Fadool et al., 2011***). For current clamp experiments, cells were held at −60mV. We sampled from the first 500ms of the current clamp experiments, before the current step began, and calculated the mean membrane potential to confirm cells were close to the target holding potential. We noted no differences between cells from Naris Occlusion and Control animals (Naris Occlusion −61.3mV ± 0.95, *n* = 20 cells and Control −58.1 ± 2.52, *n* = 23 cells, *t*-test, *t*(28) =1.11, *p* = 0.28; presented as mean ± SEM). For current step experiments, a series of 1000ms current steps were applied to evoke APs (0-500pA, 25pA steps). If multiple recordings were made from the same cell, we averaged spike counts or AP feature measurements for that cell.

#### Electrophysiology analysis

Physiology data were analyzed using custom scripts (see Source code and imaging data) written in the Python programming language (version 3.7-3.9). We used the pyABF Python module (version 2.2.8) to open axon binary format files (***Harden, 2020***). Additionally, we used the Python libraries Matplotlib 3.3.2 (***Hunter, 2007***), numpy 1.19.2 (***Harris et al., 2020***), and scipy 1.5.2 ***(Virtanen et al., 2020***).

### Reagents and antibodies

See Table 9. The research resource identifier (RRID), chemical abstracts number (CAS), national drug code (NDC), or catalog number (CAT) for the chemical, drug, or antibody are given. Not applicable (NA) is given if this information is not available or is custom made.

**Table 9.** Reagent and antibody information.

| Reagent/antibody | Type | Manufacturer | CAT/CAS/RRID # |
| --- | --- | --- | --- |
| DNQX | Chemical/Drug | Tocris | CAT: 0189 |
| APV | Chemical/Drug | Tocris | CAT: 0105 |
| Gabazine | Chemical/Drug | Sigma | CAT: S106 |
| K-gluconate | Chemical/Drug | Sigma | CAT: G4500 |
| Glucose | Chemical/Drug | Sigma | CAT: G8270 |
| HEPES | Chemical/Drug | Fisher | CAT: BP310 |
| Thiourea | Chemical/Drug | Alfa Aesar | CAS: 62-56-6 |
| Na-pyruvate | Chemical/Drug | Sigma | CAT: P2256 |
| Na<sub>2</sub>-phosphocreatine | Chemical/Drug | Sigma | CAT: P7936 |
| Na-ascorbate | Chemical/Drug | Sigma | CAT: A7631 |
| EGTA | Chemical/Drug | Sigma | CAT: E3889 |
| Biocytin | Chemical/Drug | Invitrogen | CAT: B1592 |
| Mg-ATP | Chemical/Drug | Sigma | CAT: A9187 |
| Na-GTP | Chemical/Drug | Sigma | CAT: G8877 |
| CaCl<sub>2</sub> | Chemical/Drug | Sigma | CAT: C8106 |
| MgSO<sub>4</sub> | Chemical/Drug | Macron Fine Chemicals | CAT: 6066-04 |
| NaHCO<sub>3</sub> | Chemical/Drug | Sigma | CAT: S5761 |
| NaH<sub>2</sub>PO<sub>4</sub> | Chemical/Drug | Sigma | CAT: S3139 |
| NaCl | Chemical/Drug | RPI | CAT: S23020 |
| KCl | Chemical/Drug | Sigma | CAT: P9541 |
| NMDG | Chemical/Drug | Acros Organics | CAT: 126845000 |
| 32% Paraformaldehyde | Chemical/Drug | EMS | CAT: 15714-S |
| 25% Glutaraldehyde | Chemical/Drug | EMS | CAT: 16220 |
| Na-Cacodylate buffer | Chemical/Drug | EMS | CAT: 11654 |
| Fatal-Plus | Chemical/Drug | Vortech Pharmaceuticals | NDC: 0298-9373 |
| OCT | Chemical/Drug | EMS | CAT: 72592 |
| Tween-20 | Chemical/Drug | Thermo Fisher | CAT: BP337 |
| Triton X-100 | Chemical/Drug | Thermo Fisher | CAT: BP151 |
| Hoechst | Chemical/Drug | Promokine | CAT: CA707-40013 |
| Fluoromount-G | Mounting media | Southern Biotech | CAT: 0100-01 |
| Streptavidin | Chemical | Jackson ImmunoResearch | AB_2337250 |
| NeuroTrace (Nissl) | Chemical | Thermo Fisher | AB_2620170 |
| AnkG | Antibody | UC Davis/NeuroMab | RRID: AB_2877524 |
| Plp (AA3) | Antibody | *Yamamura et al. (1991)* | NA |
| TH | Antibody | Sigma | RRID: AB_477569 |
| Caspr | Antibody | UC Davis/NeuroMab | RRID: AB_2877274 |
| Na<sub>v</sub>1.6 (SCN8A) | Antibody | Alomone Labs | RRID: AB_2040202 |
| Olig2 | Antibody | Millipore | RRID: AB_10807410 |
| CC1 | Antibody | Millipore | RRID: AB_2057371 |
| PDGFRα (AP5) | Antibody | BD Biosciences | RRID: AB_2737788 |
| Goat anti Rabbit (594nm) | Secondary antibody | Thermo Fisher | RRID: AB_2534079 |
| Goat anti Rat (647nm) | Secondary antibody | Thermo Fisher | RRID: AB_141778 |
| Goat anti IgG1 (488nm) | Secondary antibody | Thermo Fisher | RRID: AB_2535764 |
| Goat anti IgG1 (647nm) | Secondary antibody | Thermo Fisher | RRID: AB_2535809 |
| Goat anti IgG2a (488nm) | Secondary antibody | Thermo Fisher | RRID: AB_2535771 |
| Goat anti IgG2a (594nm) | Secondary antibody | Thermo Fisher | RRID: AB_2535774 |
| Goat anti IgG2b (647nm) | Secondary antibody | Thermo Fisher | RRID: AB_2535811 |
| Goat anti IgG2b (594nm) | Secondary antibody | Thermo Fisher | RRID: AB_2535781 |

## Acknowledgments

We would like to thank Dr. Jennifer Bourne, manager of the University of Colorado Anschutz Medical Campus Electron Microscopy Center for advising on the design of, and performing electron microscopy experiments for this publication. We would like to thank Nicole Arevalo, MA, for her expertise and extensive assistance with experiments, animal care, and animal breeding, and Katie Given, MS, for her invaluable support and technical expertise when performing experiments for this manuscript. We would also like to thank Dr. John Caldwell and Dr. Nathan Schoppa for experimental design and data analysis advice, and for reviewing and providing helpful input to the manuscript. We would also like to thank Dr. Alexandria Hughes for helpful conversations, support, and review of the manuscript.

Research reported in this publication was supported by the National Institute On Deafness And Other Communication Disorders of the National Institutes of Health under award numbers F31 DC018459 (NMG) and R01 DC000566 (DR). Support was also provided by the University of Colorado Anschutz Neuroscience training program T32 HD041697 and a Colorado Clinical and Translation Sciences Institute training fellowship TL1 TR001082 (NMG). The content does not necessarily represent the official views of the National Institutes of Health.

